# Fast and slow learning mediated by distinct climbing fiber signals

**DOI:** 10.64898/2025.12.19.695643

**Authors:** Dimitar Kostadinov, Beverley A. Clark, Michael Häusser

**Author notes:** Correspondence: DK.

## Abstract

Learning on both fast and slow timescales is required to enable us to adapt to a dynamic environment. Whether the mechanisms mediating fast and slow learning are implemented by the same, or different, circuit elements remains an important open question. Learning involving the cerebellum is known to be driven primarily by instructive climbing fiber inputs to Purkinje cells, but the dynamics of climbing fibers across learning on fast and slow timescales are not known. It is unclear if climbing fibers encode the same or different instructive signals at different stages of learning, and whether learning-related changes in climbing fiber encoding properties across timescales are driven by the same or distinct mechanisms. We addressed this problem using longitudinal 2-photon calcium imaging of climbing fiber activity across multiple cerebellar lobules as mice learned to execute a visuomotor integration task – slow learning – and then rapidly adapted to a new sensorimotor coupling – fast learning. Instructive signals are spatially segregated in trained mice: Lobule V preferentially encodes movement predictively, while Lobule simplex preferentially receives reward-related feedback. Moreover, the encoding of sensorimotor and reward-related instructive signals is dynamic on both slow and fast timescales: movement-related activity emerges over the course of training, while reward-related activity progressively diminishes as mice become experts but re-emerges specifically in Lobule simplex when rewards become scarce. Finally, while the same climbing fiber inputs can change on both timescales, the magnitude and spatial organization of slow changes are not related to fast changes. Thus, climbing fiber input can carry distinct instructive signals on different timescales – a multiplexing of teaching signals that may underlie the cerebellum’s capacity to both acquire and refine motor behaviour.

## INTRODUCTION

A key function of nervous systems is to use experience to inform future actions and decisions – the basis of learning. Learning occurs on a variety of timescales and in a variety of different ways (1, 2). Some associations – like the pairing of a neutral sensory event with an aversive outcome like a foot-shock – can be learned by humans and other animals in a single shot (3, 4), while other types of learning occur on much longer timescales (5, 6). Goal-directed motor behaviours that require high levels of coordination and precision often require persistent practice and repetition. Becoming proficient in these types of behaviours often happens in stages: novices must first learn the structure of a task before they can refine actions to expertly produce appetitive outcomes in reward-guided tasks (5, 7–10). Once these task structures are learned by experts, small alterations to task structures can be learned much more rapidly (6, 11–13). The differing requirements of learning to execute motor tasks and refining behaviour once tasks are learned suggest that these processes may rely on distinct neural substrates and require distinct forms of instruction (13, 14). However, most studies of the neural basis of motor learning focus on one of these phases – either task acquisition (7, 15, 16) or task adaptation (17–19). Thus, it remains unclear whether neural circuits utilize the same or different neural substrates for flexible learning across timescales.

Here, we address this knowledge gap in the cerebellum using longitudinal recordings of instructive climbing fiber (CF) inputs to the cerebellar cortex in mice as they learned to perform and refined their performance of a visuomotor integration task. CF inputs to Purkinje cells are instrumental for cerebellum-dependent learning (20), so understanding what CFs communicate to the cerebellum is crucial for understanding what and how the cerebellum can learn (21). Despite their importance, most studies of the encoding properties of CFs have been performed in trained animals, after learning has plateaued. Thus, it is unclear whether encoding features of CF inputs – such as sensorimotor error signals, movement-related timing signals, and reward prediction and omission signals (14, 17, 18, 22–31) – are present in task-naïve animals or are acquired during learning. Furthermore, it is not clear how changes in encoding during task acquisition are related to signals seen in trained mice during sensorimotor adaptation, when climbing fibers are known to shape behavioural learning (17–19, 32, 33).

We have developed an approach for performing 2-photon calcium imaging to record instructive CF input signals in the same PCs chronically over weeks in two cerebellar regions – Lobules V and simplex – while head-fixed mice learned to rotate a wheel to move a visual stimulus to a target location and obtain a reward. Once mice were experts, we probed fast, adaptive learning by altering the gain between object translation and wheel movement. In this way, we were able to determine how encoding of task events emerged in CF populations during initial stages of learning and whether CFs carry distinct instructive signals to the same PC populations when behaviours are refined and adapted. We find that in trained mice, CF inputs to both Lobules V and simplex signal movement initiation and reward delivery, with a preference for movement encoding in Lobule V and reward in Lobule simplex. Movement-predictive CF signals were minimal in naïve mice, emerged as mice learned to associate their actions to rewarded outcomes, and remained stable in expert mice performing adaptation. In contrast, reward-related CF signals were dynamic on both slow and fast timescales, in particular in Lobule simplex. They were progressively suppressed as mice became experts but re-emerged when rewards became scarce during gain adaptation. Across both Lobule V and simplex, the magnitude and spatial organization of slow and fast learning-related changes were distinct, suggesting that modification across these timescales may be driven through distinct processes. Together, these results demonstrate that instructive CF signals are dynamically modulated according to the local demands of learning across multiple timescales.

## RESULTS

### Learning in a visuomotor integration task on slow and fast timescales

We trained mice to perform a virtual reality-based visuomotor integration task to study how cerebellar activity patterns evolve over learning. Mice were head-fixed in front of an array of monitors and trained to use a steering wheel placed in front of their forepaws to control a virtual object (22). Our training procedure required progression through several task stages (**Figure 1a-b**; see **Methods**). In initial ‘Naïve’ sessions, mice first associated turning the steering wheel with virtual object translation and reward delivery. Most wheel turns that moved the object towards the visual midline were rewarded, and stereotyped wheel movements were not required for mice to gain reward. After mice made this association, they progressed to performing a precise visuomotor integration task in which they were required to make single wheel movements that translated the virtual object from an eccentric visual position (+45° from the visual midline) into a target area (±15° from the visual midline) and stopped it in this position for 100 ms. If they stopped the object in the target region, mice received a delayed reward 500 ms after the object stopped (400 ms after trial evaluation). If they undershot or overshot the target, no reward was given. The first sessions in which the mice performed this task version were termed ‘Learning’ sessions, while sessions in which behavioural performance had improved and plateaued were termed ‘Expert’ sessions. After mice became experts, we introduced ‘Adaptation’ sessions in which we altered the sensorimotor coupling (i.e. the gain) between object translation and wheel movement.

**Figure 1:**
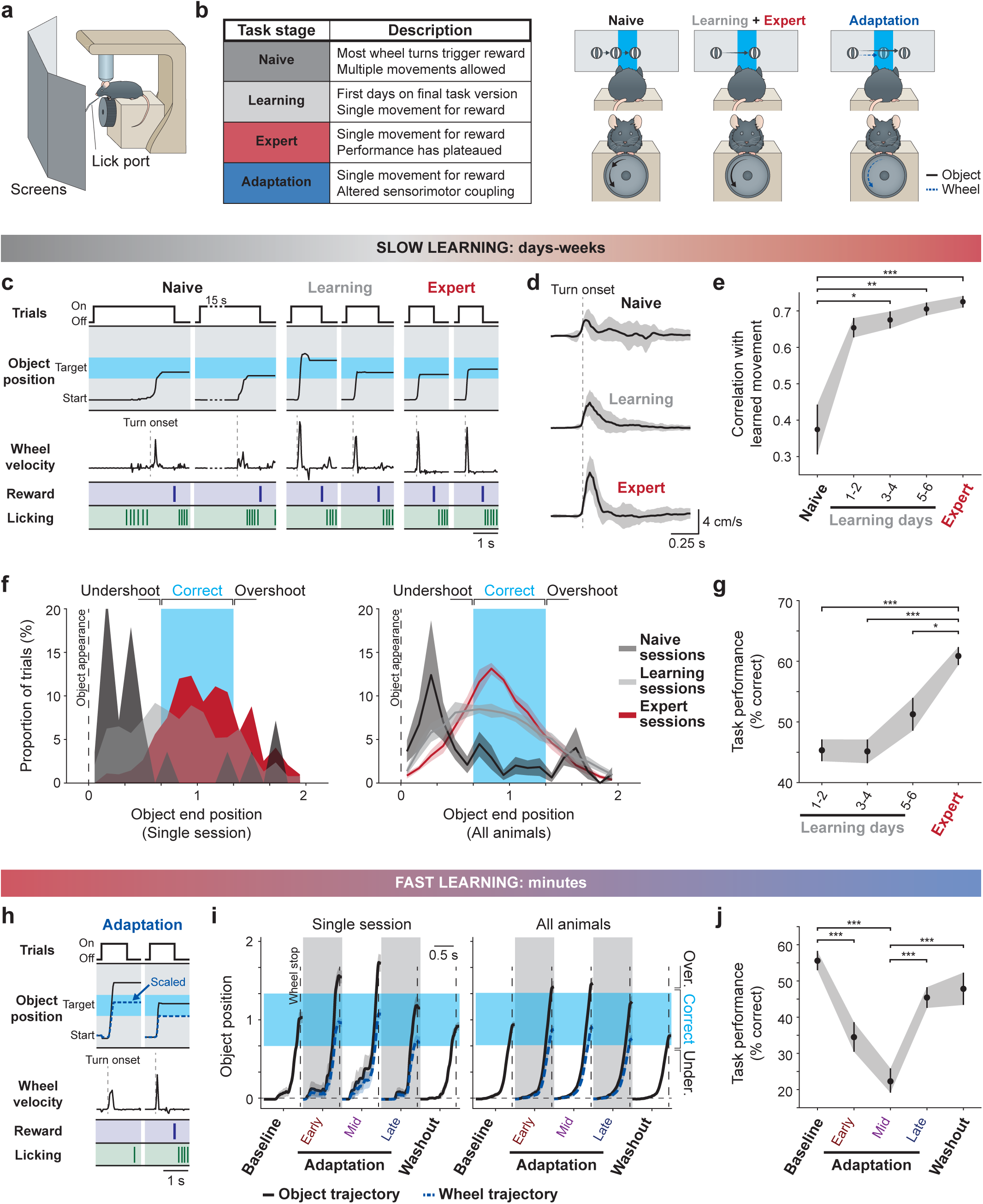
Multiple timescales of learning in a visuomotor integration task. **a.** Behavioural setup: Mice were head-fixed in front of an array of three monitors and trained to use a steering wheel with their forepaws to translate a virtual object from an eccentric visual position to the midline to obtain a delayed reward. **b. Left:** Description of the stages of the behavioural task: In ‘Naïve’ sessions, mice could make multiple wheel turns to move the object into the central target region and most trials with turns resulted in reward delivery. In ‘Learning’ sessions, mice were first required to make single wheel turns and stop the virtual object in the central target region to obtain reward. In ‘Expert’ sessions, performance on the task requiring a single continuous wheel turn for reward had plateaued. In ‘Adaptation’ sessions, sensorimotor gain changes are imposed. **Right:** schematic of the three behavioural configurations corresponding to the task stages. **c.** Example trials at different task stages showing trial start and end, object position, wheel velocity, reward delivery time, and lick times. **d.** Example wheel velocity trajectories aligned to movement onset from a single mouse at different task stages (Data are shown as mean ± s.d.; n = 73, 134, and 182 trials in ‘Naïve’, ‘Learning’, and ‘Expert’ sessions, respectively). **e.** Session-wise correlation of wheel movements at different task stages and the mean rewarded movement in ‘Expert’ sessions a.k.a. the ‘Learned movement’ (n = 9 mice). Statistics: Kruskal–Wallis test, H(4) = 17.0, P = 0.002, significance values for Bonferroni-corrected individual comparisons: Naïve versus Learning days 1-2, P = 0.1; Naïve vs Learning days 3-4, P = 0.023; Naïve vs Learning days 5-6, P = 0.006; Naïve vs Expert, P = 0.0008; all other comparisons non-significant. **f.** Probability distribution histogram of object endpoints in ‘Naïve’, ‘Learning’, and ‘Expert’ sessions for a single mouse (left) and across mice (right) (n = 9 mice). **g.** Performance on the ‘Learning’/’Expert’ task version requiring a single continuous wheel turn to translate the object into the central reward zone as a function of the number of days trained (n = 18 sessions from 9 mice). Statistics: Kruskal–Wallis test, H (3) = 28.3, P = 3 x 10^−6^, significance values for Bonferroni-corrected individual comparisons: Learning days 1-2 vs Expert, P = 2 x 10^−5^; Learning days 3-4 vs Expert, P = 2 x 10^−5^; Learning days 5-6 vs Expert, P = 0.02; all other comparisons non-significant. **h.** Example trials as in panel **c** in an ‘Adaptation’ session. **i.** Object (black) and wheel trajectories (blue) in a single session (left) and across sessions (right) for different groups of trials in the adaptation sessions: baseline trials (with normal gain), early adaptation trials (just after the gain change), mid-adaptation trials (when performance was worst), late adaptation (when performance had improved), and washout (after the change back to baseline gain) (n = 21 sessions from 9 mice). **j.** Performance on the ‘Adaptation’ task version as a function of trial groups described in panel **i** (n = 21 sessions from 9 mice. Statistics: Kruskal–Wallis test, H(4) = 41.5, P = 2 x 10^−8^, significance values for Bonferroni-corrected individual comparisons: Baseline vs Adaptation-Early, P = 5 x 10^−4^; Baseline vs Adaptation-Mid, P = 6 x 10^−8^; Adaptation-Mid vs Adaptation-Late, P = 9 x 10^−4^; Adaptation-Mid vs Washout, P = 2 x 10^−4^; all other comparisons non-significant. Data are shown as mean ± s.e.m. unless otherwise noted. Statistics summary: *\*P < 0.05, **P < 0.01, ***P < 0.001*.

Importantly, mice exhibited hallmarks of learning at each stage of the training process (6, 34). Naïve mice did not react to the object appearance consistently (**Figure 1c**). The proportion of trials with wheel turns was 49 ± 9 % in Naïve sessions and 91 ± 1 % in Learning and Expert sessions (mean ± s.e.m., n = 12 Naïve sessions and n = 72 Learning and Expert sessions from 9 mice, *P < 0.001*, Wilcoxon rank-sum test). In Naïve sessions, mice made wheel movements that were small and not stereotyped (**Figure 1d, top; Figure 1e**), and they did not produce object endpoints consistently placed at the rewarded target region (**Figure 1f**). As mice transitioned to the ‘Learning’ phase of the task, they made wheel turns that were faster, more stereotyped, and frequently resulted in the virtual object stopping in the target area (**Figure 1d-f**). These features became even more consistent as mice became ‘Experts’, and their behavioural performance improved further (**Figure 1g**). On average, ‘Naïve’ mice trained for 22 ± 2 sessions before transitioning to the ‘Learning’ stage and were trained a further 12 ± 2 sessions before their behaviour plateaued at ‘Expert’ level (mean ± s.e.m., n = 9 mice).

In a subset of the mice that became experts, we performed additional ‘Adaptation’ sessions consisting of a set of baseline trials (range 60-80), a finite number of sensorimotor gain increase trials in which well-trained movements would consistently produce overshoots of the target region if unadapted (range 120-160), and finally a set of washout trials at the original sensorimotor gain. In these ‘Adaptation’ sessions, in which learning would take place within a single hour-long training session, mice exhibited hallmarks of fast motor learning including adaptation to altered sensorimotor gain, as well as washout effects after the gain was changed back (**Figure 1h-j**).

### Longitudinal recording of climbing fiber inputs to Purkinje cells during learning

To image Purkinje cell populations during the task, we expressed the calcium indicator GCaMP in Purkinje cells by injecting Cre-dependent GCaMP6f or jGCaMP7f virus into Pcp2(L7)-cre mice. Injections were targeted to the forelimb regions of the cerebellar cortex (35, 36) hemispheric Lobule simplex and adjacent vermis lobules V and VI (**Figure 2a**), and we could readily identify calcium events in Purkinje cell dendrites, which report climbing fiber input and complex spiking with high signal-to-noise ratio (37–39) (**Figure 2b-c**). We recorded activity chronically in Lobules V and simplex as mice learned to perform the task, returning to the same fields of view across days and matching individual Purkinje cell dendrites across sessions (**Figure 2d-e**). We made recordings from both Lobules V and simplex in six mice that learned the task, recording from each region at each learning stage in at least five of six mice (**Extended Data Figure 1**). We were able to match a substantial proportion of individual PC dendrites across weeks of training and recording – 50% of dendrites were retained on average for sessions spaced by 38 days (**Figure 2f**). Importantly, the event rates of CF inputs to PC dendrites were stable across recording periods (**Figure 2g**) and the number of identified PC dendrites was not correlated with the days post injection (**Figure 2h**), indicating that changes in task encoding are unlikely to be a result of changes in indicator expression or other changes in the imaging window quality. Thus, we were able to compare activity patterns in Lobules V and simplex at each task stage and assess how activity in each region evolved during learning. In line with our previous work (22), recorded fields of view in Lobule simplex displayed the hallmarks of microzonal organization: sorting the correlation matrix of Purkinje cells orthogonally to the parasagittal axis of their dendrites revealed a block-diagonal structure that is indicative of microzones (**Extended Data Figure 2**). Our recordings in Lobule simplex capture the CF-driven activity of PC dendrites across multiple (>4) microzones. In contrast, spatially sorted correlation matrices of Lobule V fields of view often presented as one or two large blocks (despite similar numbers of PC dendrites per field of view). Thus, our recordings in Lobule V likely capture the CF-driven activity of PC dendrites from only 1-2 microzones. To make fair comparisons between Lobules V and simplex, we analyzed the encoding properties of all PC dendrites in each field of view without microzonal categorization.

**Figure 2:**
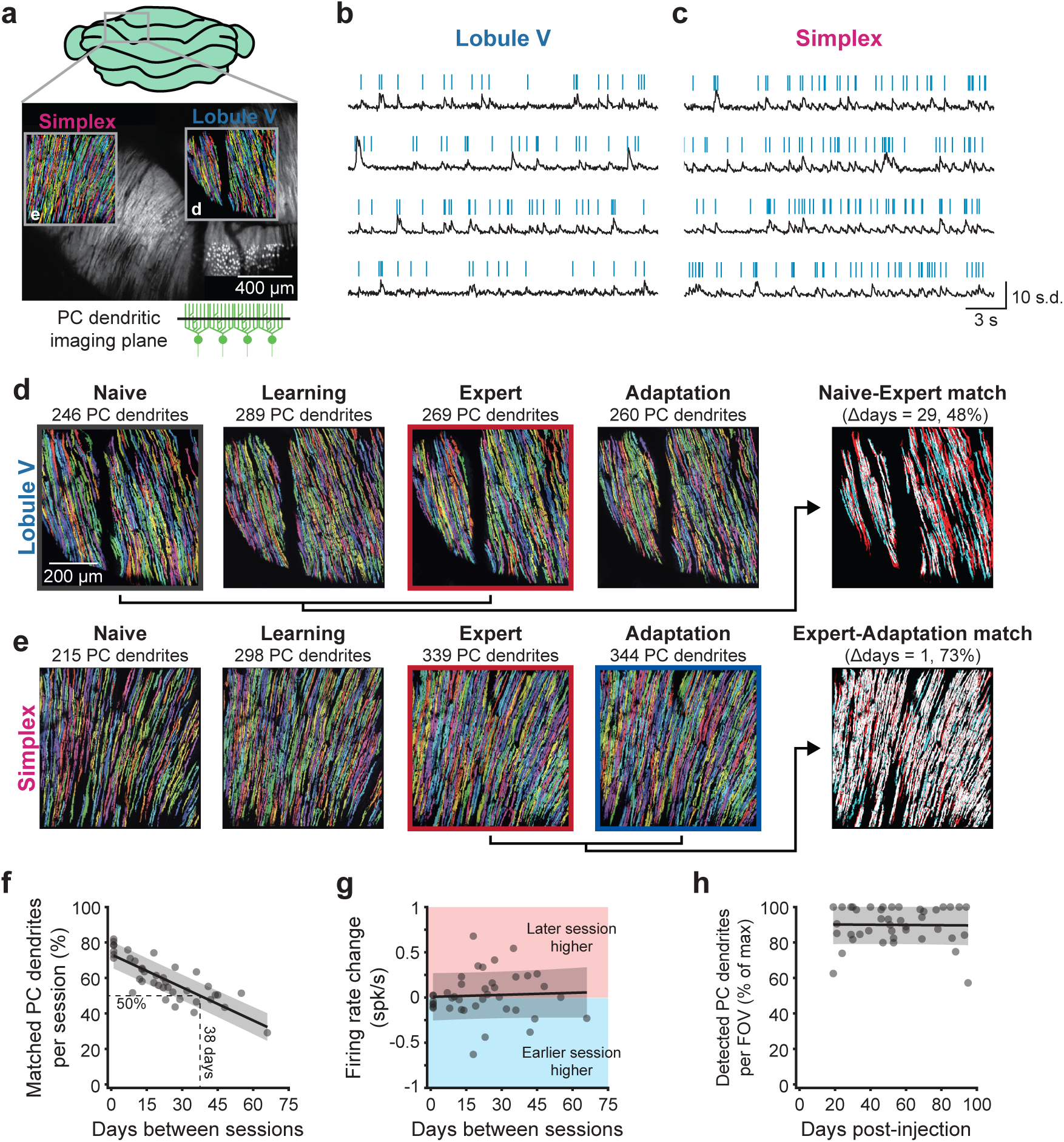
Longitudinal imaging of climbing fiber inputs to Purkinje cells during task acquisition and adaptation. **a.** GCaMP7f-labeled Purkinje cells in Lobule V and simplex. Fields of view (FOVs) correspond to those shown in panels **d** and **e**. **b.** Fluorescence traces (black) and extracted dendritic events (blue) for example Lobule V Purkinje cell dendrites. **c.** Same as panel **b** but for Lobule simplex field of view. **d.** Extracted Purkinje cell dendritic regions of interest from Lobule V field highlighted in panel **a** across ‘Naïve’, ‘Learning’, ‘Expert’, and ‘Adaptation’ sessions. Right-most panel shows PC dendrites that were matched between ‘Naïve’ and ‘Expert’ sessions (overlapping pixels of cyan and red PC dendrites shown in white). **e.** Same as panel **d** but for Lobule simplex field of view and showing matched ‘Expert’ and ‘Adaptation’ sessions. **f.** Proportion of regions of interest that could be aligned for any pair of sessions as a function of time between sessions. **g.** Detected PC dendritic event rates as a function of time between sessions. **h.** The number of PC dendrites detected in a single session as a function of days after viral injection of GCaMP6f/7f.

### Encoding properties of climbing fiber inputs to Lobules V and simplex in expert mice

We first compared the encoding properties of CF inputs to Lobules V and simplex in expert mice (**Figure 3a**), analyzing sessions from six mice where recordings were made across both lobules. Importantly, both the movement statistics (**Figure 3b**) and outcome statistics (**Figure 3d**) were indistinguishable across these sessions, and session-wise activity patterns matched our previous recordings in expert sessions of the task (22): movement-aligned activity was similar across trial outcomes (**Figure 3c**) and reward delivery evoked a strong response on correct trials (**Figure 3e**). However, task-aligned CF-driven activity of individual PC dendrites in Lobules V and simplex showed important differences (**Figure 3f**): while CF inputs to Lobule V were preferentially active immediately prior to movement onset and weakly modulated at the time of reward delivery, CF inputs to Lobule simplex were preferentially modulated at the time of reward delivery. While these results suggest that instructive signals in Lobules V and simplex may carry complementary streams of information, they are not on their own conclusive. This is because wheel movement and reward delivery often co-occurred with other task events, such as object appearance and licking.

**Figure 3:**
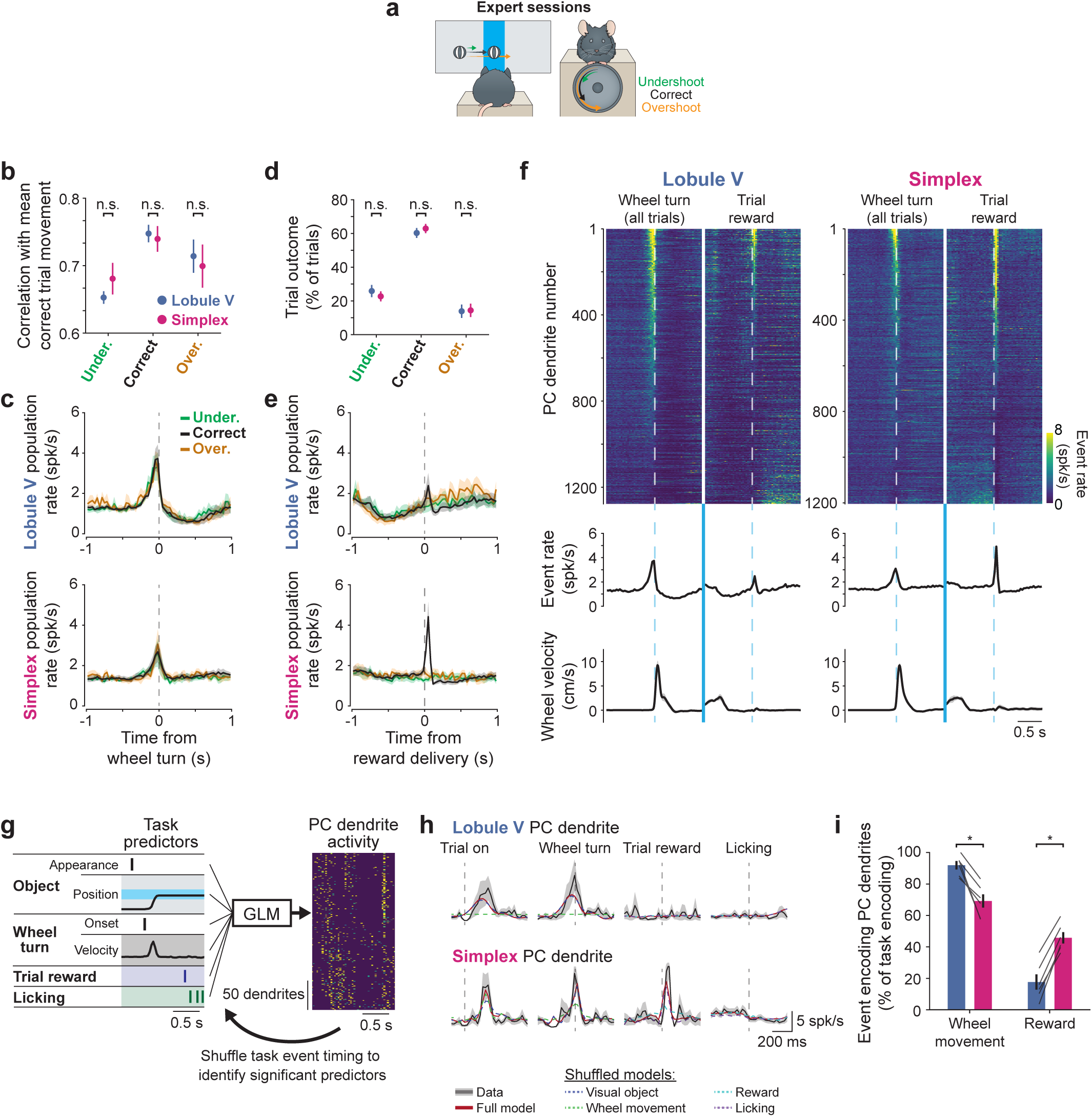
Differential encoding of movement and outcome by climbing fiber inputs to Lobules V and simplex in expert mice. **a.** Schematic of behaviour on Expert sessions, which could result in undershoots, correct trials, or overshoots. **b.** Session-wise correlation of wheel movements resulting in the three types of trial outcome and the mean correct movement a.k.a. the ‘Learned movement’ for sessions in which recordings were made in either Lobule V (blue) or simplex (magenta) (n = 6 sessions each from 6 mice, 1 session per field of view per mouse). Statistics: Lobule V vs simplex: P = 0.22 [Undershoots]; P = 0.69 [Corrects]; P = 1 [Overshoots], Wilcoxon signed-rank test. **c.** Session-wise averaged PC dendritic event rates in Lobule V (**top**) and simplex (**bottom**) aligned to wheel movement onset in undershoot, correct, and overshoot trials (n = 6 sessions each from 6 mice, 1 session per field of view per mouse). **d.** Proportion of undershoot, correct, and overshoot trials for sessions in which recordings were made in either Lobule V (blue) or simplex (magenta) (n = 6 sessions each from 6 mice, 1 session per field of view per mouse). Statistics: Lobule V vs simplex: P = 0.56 [Undershoots]; P = 0.69 [Corrects]; P = 0.69 [Overshoots], Wilcoxon signed-rank test. **e.** Same as panel **c** but aligned to expected trial reward time (500 ms after movement offset) in undershoot, correct, and overshoot trials (n = 6 sessions each from 6 mice, 1 session per field of view per mouse). **f. Top**: Trial-averaged PC dendritic event rate in Lobule V and simplex aligned to wheel movement onset (left column; pooled across trial outcomes) and trial reward (right column). PC dendrites were pooled across 6 recordings from 6 mice each and are sorted separately for each event by the first coefficient of principal component analysis (PCA) performed over the interval ±500 ms (n = 1277 Lobule V and 1204 simplex PC dendrites from 6 mice each). **Middle**: Mean time course of dendritic event rate aligned to movement onset (left) and operant reward (right) (n = 6 mice). **Bottom**: Mean time course of steering wheel velocity aligned to movement onset (left) and operant reward (right). **g.** A generalized linear model (GLM) was fit to neural activity of each PC dendrite based on task and behavioural features and used to predict activity trials held out from the fitting procedure. See **Extended Data** Figure 3. **h.** Example GLM analysis for one example PC dendrite in Lobules V and simplex each showing data (black traces), the full GLM prediction (red traces), and GLMs trained with visual object, wheel movement, reward, and licking predictors shuffled in time. Note that shuffling some but not other predictors degrades model performance. **i.** Summary of proportions of task-encoding PC dendrites with significant wheel movement and trial reward encoding in Lobule V and simplex (n = 6 sessions in 6 mice, 1 session per field of view per mouse). Statistics: Lobule V vs simplex: P = 0.031 [Wheel movement]; P = 0.031 [Reward]; Wilcoxon signed-rank test. Data are shown as mean ± s.e.m. unless otherwise noted. Statistics summary: *\*P < 0.05, n.s. = non-significant*.

We therefore disambiguated encoding of co-occurring task events by fitting generalized linear models (GLMs) in which we modeled the activity of each CF input based on task epochs and behavioural events, including trial onset (object appearance), object position, steering wheel velocity, reward delivery, and licking (**Figure 3g**) (40–44). Most CF inputs to PC dendrites in both Lobule V (92 ± 2 %, mean ± s.e.m., n = 6 fields of view from 6 mice) and simplex (80 ± 6 %, mean ± s.e.m., n = 6 fields of view from 6 mice) exhibited significant task modulation. To identify which specific task parameters were encoded by individual CF inputs, we compared the predictive power of the full model to models in which we shuffled the timing of some task parameters and not others (**Figure 3h; Extended Data Figure 3**). If shuffling a given set of predictors deteriorated the model’s predictive power significantly, this predictor set was considered to be encoded by CF input. We were able to intuitively visualize the predictive power of each model by computing task event-aligned heatmaps for each field of view and comparing heatmaps of recorded activity to the activity predicted by the full models as well as by models in which specific predictors were shuffled. For example, it was clear that shuffling forelimb movement-related predictors degraded model performance for CF inputs to PC dendrites in Lobule V and shuffling reward-related predictors degraded model performance for CF inputs to PC dendrites in Lobule simplex (**Extended Data Figure 4a-b**). Importantly, shuffling other, temporally co-varying predictors did not result in a degradation of model performance. Overall, this analysis demonstrated that while CF inputs to PC dendrites in both Lobule V and simplex could encode both forelimb movements and reward, inputs to Lobule V were more likely to encode movement and inputs to Lobule simplex were more likely to encode reward (**Figure 3i**). In Lobule V, 896 of 1085 (83%) of these task-encoding CF inputs to PC dendrites were movement-activated and 189 of 1085 (17%) were movement-suppressed. In contrast, in simplex 656 of 750 (87%) were movement-activated and 94 of 750 (13%) were movement-suppressed. Importantly, reward-related responses were bi-directional: in Lobule V 97 of 219 (44%) were reward-activated and 122 of 219 (56%) were reward-suppressed, and in simplex 189 of 449 (39%) were reward-activated and 260 of 449 (61%) were reward-suppressed. These proportions of activated and suppressed CF inputs to PC dendrites are consistent with our previous study in Lobule simplex (22) and demonstrate that we are sampling from a variety of PC modules (21). In contrast to movement and reward, visual object- and licking-related signals were similarly represented in each region (**Extended Data Figure 4c**). The likelihood to represent any combination of task or behavioural features was generally independent, except that CF inputs to PC dendrites in Lobule simplex that encoded wheel movement were less likely than chance to encode reward (**Extended Data Figure 4d**).

While many CF inputs to PC dendrites in both Lobule V and simplex exhibited movement and reward encoding, they may do so in different ways. We therefore compared the responses of movement and reward encoding CFs in Lobules V and simplex under different task conditions. We first split trials based on movement kinematics - peak displacement, velocity, and acceleration - and compared CF activity in these trial groups. This analysis showed no systematic relationship between kinematics and activity (**Extended Data Figure 5a-b**). Given the lack of kinematic encoding, we tested if activity of CF inputs to PC dendrites may vary systematically with the accuracy of the trained movement. To test this idea, we grouped trials based on their correlation with the mean rewarded (correct) movement. Splitting trials based on this ‘correctness’ parameter revealed that activity in Lobule V was larger on trials in which the mice executed movements that resembled the rewarded movement, while activity in simplex was unchanged for the most ‘correct’ movements (**Extended Data Figure 5c**).

Given the known history-dependence of reward signals in the dopamine system (45) and our prior work demonstrating that unexpected rewards evoke larger responses than expected ones (22), we tested if reward encoding in CF inputs to PC dendrites was modulated by reward history on recent task trials. We split correct trials based on previous trial outcome (correct, undershoot, overshoot). This analysis revealed that operant reward responses of CF inputs to PC dendrites in simplex were larger when the previous trial had been incorrect, especially if the incorrect trial was an overshoot (**Extended Data Figure 5d**) whereas those in Lobule V were unaffected.

Thus, the dendrites of many PCs in both Lobules V and simplex receive CF inputs that encode movement and reward but do so in different ways. Both regions exhibit movement-predictive activation but only activity levels in Lobule V PC dendrites is correlated with the accuracy of trained movements. Conversely, both regions also exhibit reward responses (though to different extents), but only activity in inputs to Lobule simplex are modulated on fast timescales by recent reward history.

### Slow learning – climbing fiber dynamics during task acquisition

To determine how CF encoding properties changed during task acquisition, we first compared task-related activity of the same neurons in Naïve, Learning, and Expert sessions. Aligning CF-driven activity to wheel turns and trial rewards revealed substantial differences in task encoding: wheel movements were weakly represented in Naïve session recordings while trial rewards evoked substantially greater activity in Naïve sessions in both Lobules V and simplex (**Figure 4a-b**).

**Figure 4:**
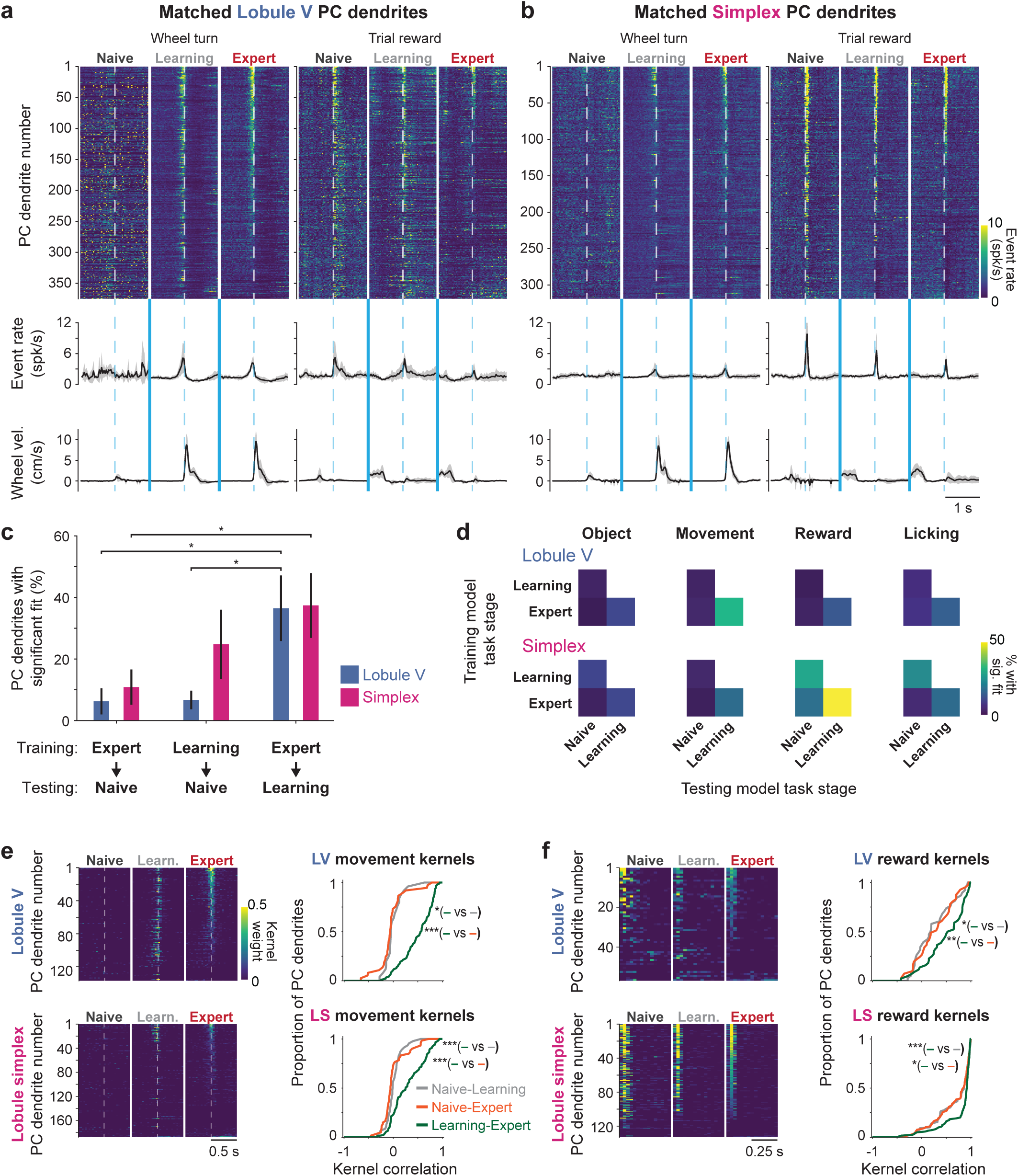
Slow learning - climbing fiber dynamics during task acquisition. **a. Top**: Trial-averaged event rates in PC dendrites matched across ‘Naïve’, ‘Learning’, and ‘Expert’ sessions in Lobule V aligned to wheel movement onset (left column; pooled across trial outcomes) and trial reward (right column). PC dendrites are sorted separately for each event by the first coefficient of principal component analysis (PCA) performed over the interval ± 500 ms (n = 365 Lobule V PC dendrites from 5 mice). **Middle**: Mean time course of dendritic event rate aligned to movement onset (left) and operant reward (right). **Bottom**: Mean time course of steering wheel velocity aligned to movement onset (left) and operant reward (right). **b.** Same as panel **a** but for Lobule simplex fields of view (n = 319 Lobule simplex PC dendrites from 5 mice). **c.** Summary of proportion of PC dendrites with significant full model fits across stages of the learning process in Lobule V (blue) and simplex (magenta). Models trained on recordings in ‘Expert’ sessions were used to predict activity recorded during ‘Naïve’ and ‘Learning’ sessions, and models trained on recordings in ‘Learning’ sessions were used to predict activity recorded during ‘Naïve’ sessions’ (n = 5 sessions in 5 mice, 1 session per field of view per mouse). Statistics: Lobule V – Friedman test, Chi-squared(2) = 7.89, P = 0.02, significance values for Bonferroni-corrected individual comparisons: Learning predicting Naïve, P = 1; Expert predicting Naïve, P = 0.04; Expert predicting learning, P = 0.04. Lobule simplex – Friedman test, Chi-squared(2) = 7.11, P = 0.03, significance values for Bonferroni-corrected individual comparisons: Learning predicting Naïve, P = 0.54; Expert predicting Naïve, P = 0.02; Expert predicting learning, P = 0.54. **d.** Summary of proportion of PC dendrites in Lobules V and simplex exhibiting significant encoding of the same task parameters across stages of learning, computed as a proportion of PC dendrites that exhibited significant task encoding using the full model fit (n = 5 sessions in 5 mice, 1 session per field of view per mouse). Rows correspond to model training stage, columns correspond to model testing stage, colorbar represents percent of CF inputs to PC dendrites with significant fit for the given task feature. **e. Left:** Movement-specific kernels fit in ‘Naïve’, ‘Learning’, and ‘Expert’ sessions independently in Lobules V (top) and simplex (bottom) for PC dendrites identified as movement-encoding using GLM analysis in Expert sessions. **Right:** Cumulative histograms of correlations between movement kernels at different task stages. Statistics: Lobule V – Friedman test, Chi-squared(2) = 19.00, P = 7.5 x 10^−5^, significance values for Bonferroni-corrected individual comparisons: Naïve-Learning vs Naïve-Expert, P = 0.25; Naïve-Learning vs Learning-Expert, P = 0.03; Naïve-Expert vs Learning-Expert, P = 4 x 10^−5^. Lobule simplex – Friedman test, Chi-squared(2) = 23.75, P = 7 x 10^−6^, significance values for Bonferroni-corrected individual comparisons: Naïve-Learning vs Naïve-Expert, P = 1; Naïve-Learning vs Learning-Expert, P = 2.7 x 10^−4^; Naïve-Expert vs Learning-Expert, P = 2.3 x 10^−5^. **f.** Same as panel **e** but for reward-specific kernels. Statistics: Lobule V – Friedman test, Chi-squared(2) = 11.03, P = 0.004, significance values for Bonferroni-corrected individual comparisons: Naïve-Learning vs Naïve-Expert, P = 1; Naïve-Learning vs Learning-Expert, P = 0.02; Naïve-Expert vs Learning-Expert, P = 0.008. Lobule simplex – Friedman test, Chi-squared(2) = 14, P = 9 x 10^−4^, significance values for Bonferroni-corrected individual comparisons: Naïve-Learning vs Naïve-Expert, P = 1; Naïve-Learning vs Learning-Expert, P = 9.9 x 10^−4^; Naïve-Expert vs Learning-Expert, P = 0.02. Data are shown as mean ± s.e.m. unless otherwise noted. Statistics summary: *\*P < 0.05, **P < 0.01, ***P < 0.001*.

Consistent with these qualitative differences, GLMs trained on Learning or Expert sessions predicted task-related activity in Naïve sessions poorly, while GLMs trained on Expert sessions exhibited significantly greater predictive power in Learning sessions (**Figure 4c**). This observation held at the level of encoding of individual task features: most PC dendrites that exhibited significant movement encoding in Learning and Expert sessions did not do so in Naïve sessions, while many reward-coding CF inputs to PC dendrites identified in Learning and Expert sessions did exhibit this encoding in Naïve sessions (**Figure 4d**).

To understand the evolution of task-related CF activity across learning in more detail, we examined the mapping between neural activity and either wheel movements or rewards at different task stages. To do this, we (1) fit GLMs with a given set of predictors shuffled in time (as in **Figure 3g**), (2) subtracted this fit from the recorded activity, yielding an activity residual, and (3) refit a model to the residual activity using only the predictors of interest. This procedure allowed us to determine the time-varying activity kernels associated with different task events and compare them across task stages. These analyses revealed that movement kernels emerged in Learning sessions, and that these were more similar between Learning and Expert sessions than between Naïve and either Learning or Expert sessions, especially in Lobule V (**Figure 4e**). Conversely, reward kernels were present even in naïve animals and remained relatively consistent across learning, especially in Lobule simplex (**Figure 4f**).

We observed that changes in task encoding across learning were more pronounced in Lobule V than in simplex. To test whether changes in task encoding may reflect a more prominent re-organization of the correlation structure of climbing fiber inputs (22, 46), we compared the similarity of correlation matrix structure of CF inputs to PC dendrites across learning in Lobule V and simplex (**Extended Data Figure 6**). The correlation matrix structure in both Lobule V and simplex was recognizably similar across Naïve and Expert sessions (**Extended Data Figure 6a-b**). Moreover, this structure was highly non-random: shifting spatially sorted PC dendrites by a single index – i.e. comparing the correlation of neurons to their nearest neighbors – resulted in a marked drop in correlation matrix similarity and completely shuffling these indices resulted in near-zero similarity (**Extended Data Figure 6c**). Across both Lobule V and simplex, the similarity of correlation matrices dropped off as a function of days between recording sessions. However, this drop-off was more pronounced in Lobule V than in simplex (**Extended Data Figure 6d**). Thus, the greater stability in task encoding observed in Lobule simplex during task acquisition is mirrored by a more stable correlation structure of climbing fiber inputs to PC populations.

Do nearby Purkinje cells exhibit similar learning-related changes in their encoding properties – as would be expected if learning-related changes show modular organization? To answer this question, we compared the similarity of kernel correlations across Naïve, Learning, and Expert task stages (**Figure 4e-f**) as a function of inter-dendritic distance. This analysis demonstrated that the degree to which nearby PC dendrites (those <100 μm apart) change across learning is more similar when compared with PC dendrites that are further (those ≥100 μm apart; **Extended Data Figure 7a-b**). Thus, it is likely that learning-related changes in encoding on slow timescales are organized at the level of cerebellar modules.

### Fast learning – dynamics of climbing fiber inputs during sensorimotor adaptation

We next probed how CF encoding properties may change in Adaptation sessions, in which animals must systematically alter their behaviour over the course of minutes to maintain successful task execution following a change in the gain between wheel movement and object translation (a gain increase; **Figure 5a**). We analyzed the responses of movement-encoding and reward-encoding CF inputs to PC dendrites (as identified in Expert sessions, **Figure 3**) first during the baseline trials before we imposed the gain change, then in batches of trials during the gain change, and finally in “washout trials” after the gain was restored to the trained value from Expert sessions.

**Figure 5:**
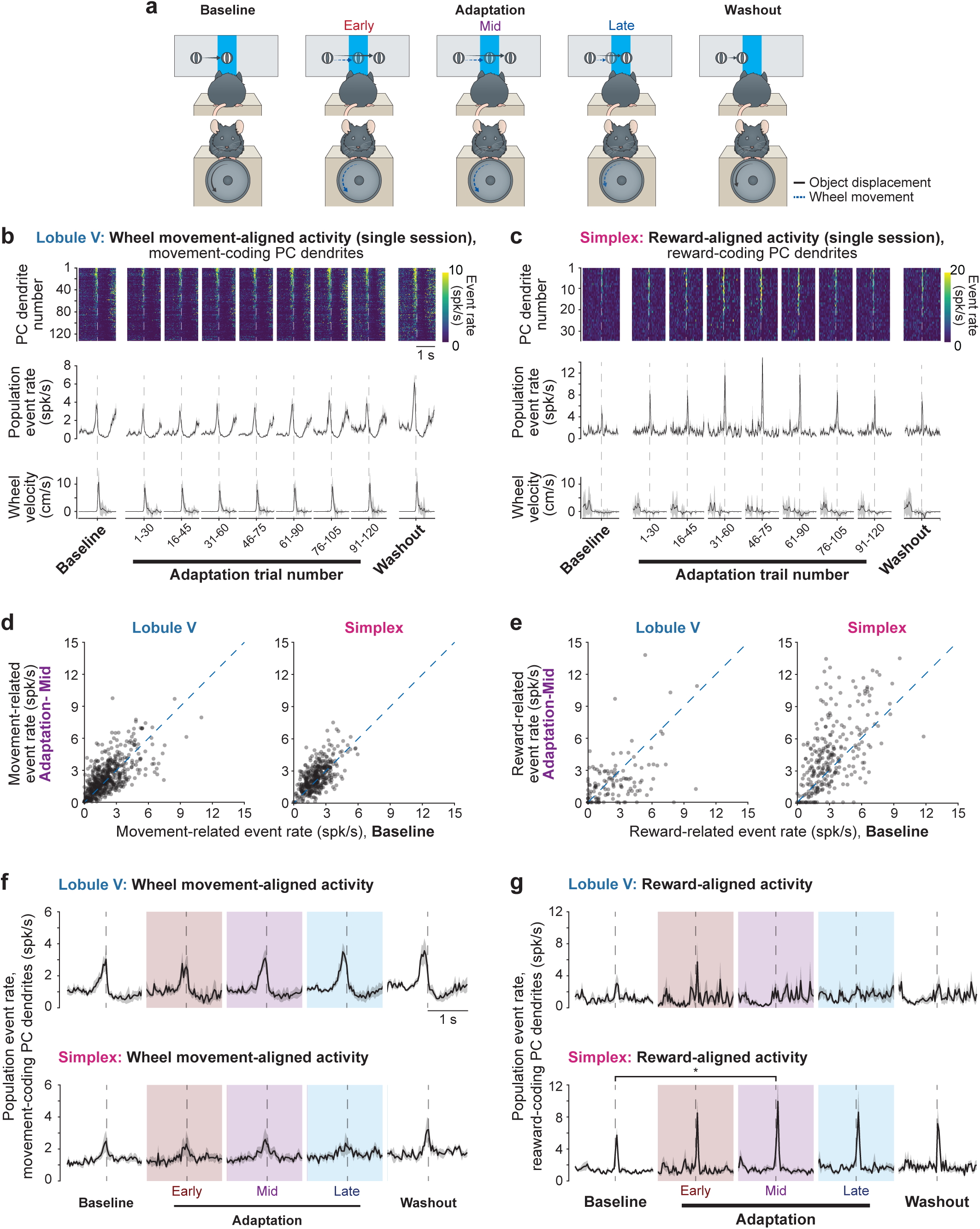
Fast learning - climbing fiber dynamics during sensorimotor adaptation. **a.** Schematic of behaviour in Adaptation sessions: baseline trials (with normal gain), early adaptation trials (just after the gain change), mid-adaptation trials (when performance was worst), late adaptation (when performance had improved), and washout (after the change back to baseline gain). **b.** Movement-aligned activity of movement-encoding PC dendrites in Lobule V of an example mouse in a single Adaptation session. Data are shown in groups of 30 trials that overlap by 15 trials. **Top**: Trial-averaged PC dendritic event rate aligned to wheel movement onset (pooled across trial outcomes) and split by trial number. PC dendrites are sorted by response magnitude (n = 132). **Middle:** Mean time course of dendritic event rate aligned to movement onset. **Bottom:** Mean time course of steering wheel velocity aligned to movement onset. **c.** Reward-aligned activity of reward-encoding PC dendrites in Lobule simplex of an example mouse in a single Adaptation session. Data are shown in groups of 30 trials that overlap by 15 trials. **Top**: Trial-averaged PC dendritic event rate in an example Lobule simplex recording session aligned to trial reward and split by trial number. PC dendrites are sorted by response magnitude (n = 34). **Middle:** Mean time course of dendritic event rate aligned to trial reward. **Bottom:** Mean time course of steering wheel velocity aligned to trial reward. **d.** Scatter plots showing pairwise comparisons of movement-related activity in movement-encoding PC dendrites in Lobule V (left, n = 602 dendrites from 5 mice) and Lobule simplex (right, n = 413 dendrites from 5 mice) during baseline trials and mid-adaptation trials (computed as mean activity over −300 to 0 ms from movement onset). **e.** Scatter plots showing pairwise comparisons of reward-related activity in reward-encoding PC dendrites in Lobule V (left, n = 128 dendrites from 5 mice) and Lobule simplex (right, n = 274 dendrites from 5 mice) during baseline trials and mid-adaptation trials (computed as mean activity over 0 to +200 ms from reward delivery). **f.** Average movement-aligned activity in Lobule V (top) and simplex (bottom) for baseline, adaptation (early, mid, and late), and washout trials (n = 5 sessions in 5 mice, 1 session per field of view per mouse). Statistics: Lobule V – Friedman test, Chi-squared(4) = 9.12, P = 0.0582 (non-significant); Lobule simplex – Friedman test, Chi-squared(4) = 3.68, P = 0.451 (non-significant). **g.** Average reward-aligned activity in Lobule V (top) and simplex (bottom) for baseline, adaptation (early, mid, and late), and washout trials (n = 5 sessions in 5 mice, 1 session per field of view per mouse). Statistics: Lobule V – Friedman test, Chi-squared(4) = 1.09, P = 0.896 (non-significant); Lobule simplex – Friedman test, Chi-squared(4) = 13.92, P = 0.0076, significance values for Bonferroni-corrected individual comparisons: baseline vs mid-adaptation, P = 0.027. Data are shown as mean ± s.e.m. Statistics summary: *\*P < 0.05*.

Movement-related activity in Lobule V and simplex did not change during adaptation. Response magnitudes and time-courses were similar across baseline, adaptation, and washout trials in individual neurons within a field of view (**Figure 5b**). Moreover, a scatter plot of movement-related activity in all identified movement-encoding CF inputs to PC dendrites comparing baseline and mid-adaptation trials showed that responses were clustered near the unity line in both cerebellar regions (**Figure 5d**). Finally, the average population movement-related response in baseline, adaptation, and washout trials across mice showed no difference across trial groupings (**Figure 5f**).

On the other hand, reward-related activity in Lobule simplex was highly dynamic during adaptation. Reward-related signals in simplex increased gradually over the dynamic phase of adaptation before decreasing in magnitude as mice stabilized their movement in response to the new condition (shown for an example session in **Figure 5c**). These population-level changes reflect scaling of responses of individual neurons, rather than the emergence of a distinct population that encodes reward specifically during adaptation, since reward-related event rates during baseline trials were correlated with reward scaling during adaptation (**Figure 5e**). These effects were also prevalent when comparing the average reward-related activity across mice – reward responses mid-adaptation were significantly larger than baseline trials before scaling down during late-adaptation and washout (**Figure 5g**).

In contrast to the scaling of reward-related signals on correct trials during adaptation, activity on incorrect trials (undershoots and overshoots) was unchanged relative to baseline trials on Adaptation sessions. Specifically, in both Lobules V and simplex, activity in movement-coding as well as reward-coding CF inputs to PC dendrites was similar around the time that the reward would have been given for both undershoot and overshoot trials (**Extended Data Figure 8**).

Thus, CF inputs to PC dendrites are also dynamic on fast timescales over minutes in a specific manner: reward-related activity in Lobule simplex grows transiently during the dynamic phase of adaptation.

### Relationship between dynamics of learning signals on slow and fast timescales

Finally, we set out to determine the relationship between the learning-related changes we had observed on slow and fast timescales. Specifically, we probed whether the changes in CF inputs to PC dendritic signals over slow timescales were related to the extent to which they change on fast timescales – suggesting that they are driven by a common process – or whether slow and fast learning-related changes were not predictive of each other – suggesting that they are distinct (**Figure 6a**). We identified PC dendrites that were recorded across Naïve, Expert, and Adaptation sessions that exhibited encoding of movement and reward and compared the extent to which their learning-related changes during task acquisition resembled those during adaptation. We reasoned that if slow and fast learning changes have common drivers, CF inputs to PC dendrites with low kernel correlations between naïve and expert mice should exhibit strong adaptation scaling (and vice-versa) – i.e. these should be negatively correlated. Alternatively, if they are driven by distinct and mutually exclusive processes, slow learning kernel correlations should be positively correlated to adaptation scaling. Finally, if they are driven by distinct but non-interacting processes, kernel correlations and adaptation scaling should be uncorrelated (**Figure 6b**). This analysis demonstrated that the degree of learning-related changes in movement-coding and reward-coding CF inputs to PC dendrites in both Lobules V and simplex are not correlated (**Figure 6c-d**). These distinct learning-related changes are consistent with the different spatial arrangement of CF inputs to PC dendrites that changed on slow and fast timescales. While CF inputs to nearby PC dendrites (those <100 μm apart) are more likely to exhibit similar slow learning kernel correlations than those further apart (those ≥100 μm apart) (**Extended Data Figure 7a-b**), adaptation-related changes in CF inputs to PC dendrites did not exhibit any clear spatial clustering in their degree of adaptation, nor in their likelihood to scale in the same direction (**Extended Data Figure 7c-d**). Thus, slow (across days-weeks) and fast (across minutes) learning-related changes in CF inputs to PC dendrites are likely driven by distinct, non-interacting processes.

**Figure 6:**
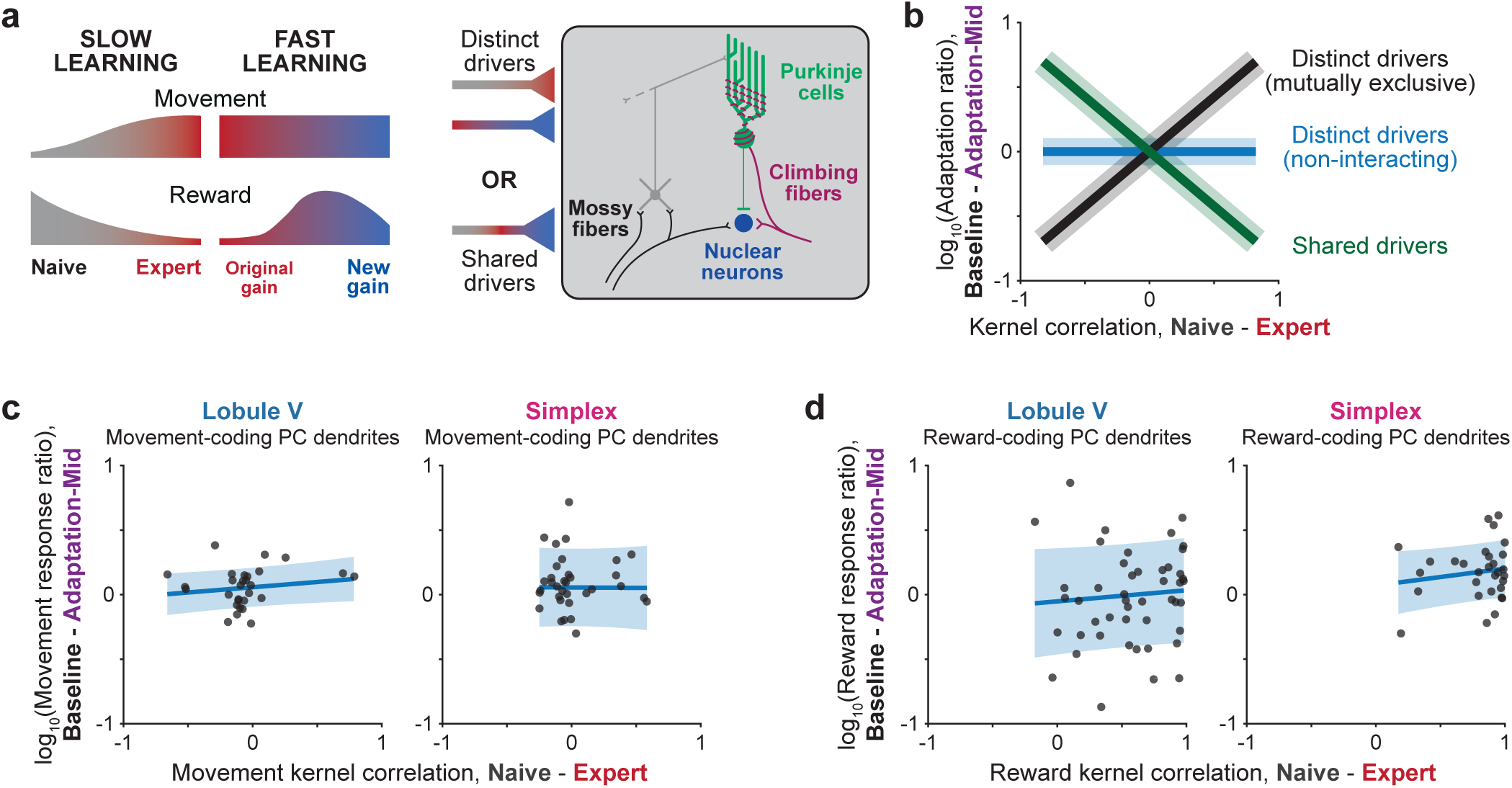
Relationship between dynamics of learning signals on slow and fast timescales. **a.** Schematic depicting shared and distinct inputs driving slow and fast learning. **b.** Potential analysis outcomes: If fast and slow learning are mechanistically related, then Naïve-Expert task kernel correlations should be anti-correlated to response ratios on Adaptation sessions (green). If these are distinct and non-interacting processes, slow and fast learning parameters should be uncorrelated (blue). Finally, if they are distinct but mutually exclusive processes, slow and fast learning parameters should be positively correlated (black). **c.** Scatter plot and linear fit (with standard error estimate) of movement kernel correlations (comparing Naïve to Expert sessions) and movement response ratios (comparing baseline to mid-adaptation responses on a log scale) in movement-encoding PC dendrites in Lobule V (n = 29, R^2^ = 0.03, P = 0.41) and simplex (n = 35, R^2^ = 2 x 10^−5^, P = 0.98).

## DISCUSSION

Climbing fiber inputs to Purkinje cells are the key drivers of cerebellum-dependent learning (20, 47). Most studies of the encoding properties of these inputs in behaving animals have investigated their properties in trained animals (18, 25–28, 32, 35, 39, 48–51). We set out to identify whether and how the encoding properties of inputs to individual Purkinje cells are shaped by learning by imaging task-related activity longitudinally in the same neuronal populations across learning over weeks. We have quantitatively characterized the encoding properties of climbing fiber inputs to Purkinje cells in cerebellar Lobules V and simplex, regions known to be involved in forelimb motor control (36) that receive input via globally similar but locally heterogeneous long-range connections with the forebrain (52), in the context of a forelimb visuomotor integration task. In trained mice, Lobules V and simplex exhibit complementary encoding of movement and reward. These signals are shaped by learning as animals become task-proficient: movement-predictive activation emerges during task acquisition while reward responses diminish. During Adaptation sessions in trained mice, movement-predictive activity remains stable while reward-related activation scales transiently in Lobule simplex. These two types of learning-related changes are not correlated: the degree of response scaling during task acquisition is not predictive of the degree of reward-related scaling, and these processes exhibit distinct spatial organization. Thus, cerebellar instructive signals conveyed by climbing fibers are dynamically shaped by learning on multiple timescales through distinct processes.

### Differential encoding of movement and reward in Lobules V and simplex of trained mice

The distribution of climbing fiber inputs to cerebellar Purkinje cells is highly organized, with specific regions of the inferior olive projecting to specific locations in the cerebellar cortex (53). This anatomical organization provides the substrate for functional organization of the cerebellum into ‘regions’ controlling forelimbs (36), whiskers (54), saccadic eye movements and compensatory visual reflexes (55, 56), licking (30), conditioned eye blinks (57), and many others. However, cerebellar modules within these regions can be heterogeneous and exhibit heterogeneous functional tuning (58). We quantitatively categorized the encoding properties of climbing fiber inputs within the forelimb region of the cerebellar cortex using GLMs. Inputs to Lobule V and simplex exhibit complementary encoding: while both regions receive both movement and reward-related information, climbing fiber inputs to Lobule V preferentially convey movement-predictive signals and inputs to simplex preferentially convey reward-related feedback signals. Thus, these regions may contribute to task execution in complementary ways by shaping outgoing motor commands (59, 60) in real time (Lobule V) and signaling trial outcome (Lobule simplex), respectively. The result that movement-related signals in Lobule V scale with movement accuracy but not with movement kinematics (**Extended Data Figure 5**) is particularly interesting, as it implies that the CF input to Lobule V is not a generic motor efference copy, but is selectively tuned to the *trained* motor program (61).

### Evolution of encoding properties during task acquisition

To determine how climbing fiber encoding properties change during task acquisition, we matched individual PC dendrites across sessions and used GLMs trained on CF inputs recorded in Expert sessions to predict neural activity in naïve and partially trained (learning) mice. GLMs trained on Expert session data predicted activity in Naïve sessions poorly, suggesting substantially different task encoding in naïve and trained mice. Specifically, these analyses revealed two key features of learning-related changes in climbing fiber inputs. Firstly, movement-related activity emerges between Naïve and Learning sessions, during which the relationship between the execution of the task and reward is established. These learned signals likely reflect a task context-specific signal (62) that would not interfere with other movement-related activity in Lobules V and simplex regions, such as that associated with locomotion (63–65). Secondly, reward-related activity is present from Naïve sessions onward but is sculpted by learning to suppress responses when rewards are more expected (22, 28, 35). In summary, task-related climbing fiber signals are deployed flexibly: predictive signals emerge during task acquisition while responses to predictable events are suppressed.

### Scaling of reward-related feedback during task adaptation

Climbing fiber inputs to PCs were also dynamic on fast timescales in expert mice challenged to perform sensorimotor adaptation, but these signals were confined to specific task events in specific cerebellar subregions. Movement-related activity in both Lobule V and simplex was stable across Adaptation sessions. On the other hand, reward-related activity, specifically in Lobule simplex, was highly dynamic during adaptation: these signals were larger during the dynamic phase of adaptation and became smaller once animals adjusted to the new sensorimotor gain. When we restored the original, well-trained, sensorimotor gain at the end of the sessions (washout period), reward responses remained slightly above baseline. These adaptation-related dynamics were confined to reward-related responses. Overshoot error trials, which became common during adaptation due to the direction of the gain change, did not alter feedback-related signals in movement- or reward-coding climbing fiber inputs to either Lobule V or simplex. Thus, reward-related feedback in lateral cerebellar regions may represent a specific pathway for tuning behavioural performance on short timescales. These results are consistent with our own previous work (22), as well as several other studies (35, 59). However, we cannot rule out that, in task designs in which feedback signals on adaptation trials may drive stronger feedback responses, error-related climbing fiber responses may be more abundant (19, 60).

### Relationship between slow and fast learning

Are the changes associated with learning on slow and fast timescales related to each other? If they are, we would expect that (1) the degree of learning-related changes during task acquisition should be related to the degree of learning-related changes during task adaptation, and (2) the relationship between learning-related changes in neighboring PC dendrites should be organized similarly. We tested these predictions directly by comparing slow and fast learning metrics to each other and measuring the similarity of learning-related changes as a function of inter-PC distance. The degree of slow learning-related changes, as measured by the Naïve-Expert session GLM kernel correlations, were uncorrelated to fast learning-related changes, as measured by the scaling of movement and reward responses during adaptation. Moreover, the spatial organization of slow and fast learning is distinct. The degree of slow timescale learning-related changes is more similar for closely spaced PC dendrites than for distant ones. This is consistent with slow learning being organized at the level of cerebellar modules, potentially through cerebellar cortico-nuclear-olivary feedback. On the other hand, fast learning-related scaling of reward responses during sensorimotor adaptation does not exhibit spatial selectivity, suggesting that this process is not organized through cerebellar modules. Thus, slow and fast learning-related changes in task encoding may be mediated through distinct circuit mechanisms.

### Outlook

This work opens several important avenues for future research. Within the cerebellum, a key to understanding the consequences of the dynamism of CF instructive signals will be to chronically identify and monitor corresponding changes in Purkinje cell simple spiking (66–68) across learning. Such recordings would reveal how the cerebellum informs, and is informed by, learning signals in the rest of the brain including cortex and the basal ganglia (14, 21, 52, 69). In a parallel study, we have determined the long-range connectivity of Lobules V and simplex Purkinje cells with the forebrain (52). Lobules V and simplex Purkinje cells receive monosynaptic climbing fiber inputs from segregated subregions of the inferior olive. However, these locally segregated afferents are innervated by a common set of regions in the forebrain. Within these forebrain partner regions, neurons coupled to Lobule V and simplex are spatially distinct. Thus, instructive signals to these regions arise from distinct subregions within largely overlapping forebrain ‘parent’ regions. This dynamic input to the inferior olive may be a key feature allowing cerebellar instructive signals to emerge and disappear, tuning instructive signals to task demands locally rather than maintaining predetermined and rigid heuristics. In this way, multiregional brain networks could employ flexible learning to guide the execution of complex goal-directed behaviour (70).

More generally, employing distinct mechanisms to guide learning across timescales may be an efficient way to maintain robust continual learning throughout the brain, preventing the erasure of previous learning while maintaining the capacity to flexibly learn new information. Mitigating this erasure – termed catastrophic forgetting – is a major challenge in both neuroscience and machine learning (71–73). Thus, our findings may represent a generalizable strategy for learning throughout the brain and may be useful for informing the design of robust and efficient learning algorithms in the next generation of artificial neural networks.

## DATA AVAILABILITY

The data that support the findings of this study are available from the corresponding authors upon reasonable request.

## CODE AVAILABILITY

The custom analysis code used in this study is available from the corresponding authors upon reasonable request.

## ACKNOWLEDGEMENTS

We thank Soyon Chun and Caroline Reuter for technical support. We are grateful to Arnd Roth, Maxime Beau, L. Federico Rossi, and Sam Clothier for helpful discussions. This work was supported by the Wellcome Trust (M.H.: PRF 201225/Z/16/Z and 224668/Z/21/Z; D.K.: Career Development Award 225951/Z/22/Z), ERC (M.H., AdG 695709) and EMBO (D.K., ALTF 914-2015).

## AUTHOR CONTRIBUTIONS

D.K., B.A.C., and M.H. conceived the project and wrote the manuscript. D.K. performed all experiments and analysis and made all figures.

## REPORTING SUMMARY

Further information on research design is available in the Life Sciences Reporting Summary linked to this article.

## METHODS

### Animals

All animal procedures were approved by the local Animal Welfare and Ethical Review Board at University College London and performed under license from the UK Home Office in accordance with the Animals (Scientific Procedures) Act 1986. We used male Pcp2(L7)-Cre mice (line Jdhu – B6.Cg-Tg(Pcp2-Cre)3555Jdhu/J) (74) aged between 3 and 6 months. Male mice were preferred in our task because they were larger and more willing to initiate wheel movements at the beginning of training, facilitating more rapid learning. Mice were group housed and maintained on a 12:12 day-night cycle. Behaviour was characterized in 9 mice, and 6 of these mice were used for chronic imaging experiments.

### Surgical procedures

#### Headplating, virus injection, and chronic window installation for behavioural experiments

Surgical procedures were similar to those described in Kostadinov et al., 2019 (22) with some minor modifications. A minimum of 2 hours before surgery, mice were injected with dexamethasone to reduce swelling during surgery. A single procedure was performed on each mouse lasting approximately 2 hours to install a headplate, inject virus, and install a chronic window. Mice were maintained under 1.5-2% isoflurane anesthesia, and Buprenorphine (1 mg/kg) was administered peri-operatively for analgesia. Once mice were anesthetized, headplates with an oval inner opening 7 mm long and 9 mm wide were installed over the left cerebellar cortex of each mouse (Lobule simplex and adjacent paravermis Lobules V and VI) and secured with dental cement. The headplates were centered at approximately 2 mm lateral and 2 mm caudal from lambda (just in front of the posterior tip of the interparietal bone).

After headplate implantation, we performed a 4-mm circular craniotomy, centered in the middle of the headplate hole, to expose the cerebellar cortex for virus injection and window installation. We then injected Cre-dependent GCaMP6f (75) virus (AAV1.CAG.Flex.GCaMP6f.WPRE.SV40) diluted to ∼5 x 10^12^ GC/ml or GCaMP7f (76) virus (AAV1.CAG.Flex.jGCaMP7f.WPRE.SV40) diluted to ∼2 x 10^12^ GC/ml from stock titer in 3-5 locations spanning Lobules V and VI in the paravermis and intermediate Lobule simplex. At each location, 100 nl of virus solution was pressure-injected at depths of 500, 375, and 250 µm below the cerebellar surface at 2-minute intervals. We waited ∼5 minutes after the final injection in a set of 3 injections before retracting the injection pipette. In total, ∼1-1.5 µl of diluted virus was injected per mouse. Finally, a 4-mm single-paned coverslip was press-fit into the craniotomy at a 12.5 degree lateral angle, sealed to the skull by small application of cyanoacrylate (VetBond) and fixed in place by dental cement. The conical portion of a nitrile rubber seal was then glued to the headplate with dental cement and filled with Kwik-Cast to protect the window preparation during recovery and between recording sessions. Mice were allowed to recover for a minimum of 7 days before beginning water restriction, during which time they were given post-operative analgesia as needed.

After mice had recovered from surgery, they were placed under water restriction for at least 5 days during which time they were acclimated to the recording setup and expression-checked. All mice were maintained at 80-85% of their initial weight over the course of recording experiments. Trained mice typically received all their water for the day from rewards during the behavioral task, while naïve mice were supplemented to 1 g water per day with Hydrogel.

### Behavior

#### Apparatus

A detailed description of the behavioral apparatus and training procedures was provided in a prior study (22). Briefly, mice were head-fixed in front of an array of monitors and trained to rotate a rubber Lego wheel with their forepaws from an eccentric visual position to the visual midline to receive sugar water rewards. The virtual environment was controlled using ViRMEn (77) software in Matlab. The rotation of the steering wheel translated the virtual object (a revolving black and white beach ball) displayed on the screens during each operant motor trial. Rewards consisted of ∼3 µl of a sugar water solution (5% sucrose) and were delivered through a solenoid valve (NResearch, part number 225PNC1-21) whose click was audible to the mouse. Behavioral parameters and task-related triggers were fed back to the VR system through an Arduino and a National Instruments DAQ card (NI USB-6212).

#### Motor task training protocol

Mice were trained in several stages, which were analyzed and compared as different epochs in the learning process. Initially, mice were trained to translate the virtual object, which appeared in the middle of either the left or right screens (at +45° or −45°), towards the visual midline to receive a reward, at a high wheel gain. On the first few days of training, the virtual object drifted towards the midline and triggered an auto-reward after a long delay (30 – 60 seconds). The data included in ‘Naïve’ sessions was acquired during the first 1-2 days of these training sessions. After mice learned to make wheel turns to receive rewards, they progressed to a unilateral version of the task (left trials only) that was not automatically rewarded. During these sessions, the wheel movements of individual mice became progressively stereotyped, allowing us to set a final gain for each mouse that maximized their chances of receiving rewards in gain-fixed training sessions. In these sessions, mice were required to make single wheel movements that translated the virtual object into a target area (±15° from the visual midline) and stopped it in this position for 100 ms. The mean gain across the mice used in this study was 3.3 degrees/mm corresponding to a 13.6 mm translation of the wheel to hit the center of the target (range 2.8-4.3 degrees/mm). On the first day that mice were trained on this gain-fixed task, their performance was 40-50% and plateaued at ∼60% after about 1 week of training on this final task version (**Figure 1g**). All recordings in ‘Learning’ sessions were acquired on the first 1-2 training days of these gain-fixed sessions, and all recordings in ‘Expert’ sessions were acquired after behavior had plateaued, after at least 7 days of training on gain-fixed sessions. Expert mice were then subjected to ‘Adaptation’ sessions, which consisted of 60-80 baseline trials at the previously trained visuomotor gain, followed by 120-160 trials of a 1.6x gain increase, and a washout period at the baseline gain. A minimum of 3 Expert sessions were performed between any two adaptation sessions acquired from a single mouse to minimize potential across-day residual learning effects.

Reward delivery on correct motor trials was timed at 400 ms after trial evaluation (500 ms after the wheel stopped moving) and was followed by a short (0-2 s) timeout, while incorrect motor trials were followed by a long (5-7 s) timeout. After completion of the timeout, a variable withhold period (1.5-2.5 s) was enforced, during which mice were obligated to not lick or turn the steering wheel. Licks were detected using an electrical lick circuit (78). In addition to operant rewards, mice received random or tone-cued rewards that were administered upon the completion of withhold periods. Tone cues for reward trials consisted of a 100 ms long, 4 kHz tone followed by 400 ms of silence before reward delivery.

### Data acquisition

#### Two-photon calcium imaging during behavior

Calcium imaging experiments were performed using a 16x/0.8 NA objective (Nikon) mounted on a Sutter MOM microscope equipped with the Resonant Scan box module. A Ti:Sapphire laser tuned to 930 nm (Mai Tai, Spectra Physics) was raster-scanned using a resonant scanning galvanometer (8 kHz, Cambridge Technologies) and images were collected at 512 x 512 pixel resolution over fields of view of 670 µm x 670 µm at 30 Hz. Sample plane power used for recordings ranged from 15-70 mW and recordings were performed midway between the pial surface and the Purkinje cell body layer, at depths between 40 and 100 µm. The microscope was controlled using ScanImage (Versions 2015 and 2018, Vidrio Technologies (79)) and tilted to ∼12.5 degrees such that the objective was orthogonal to the surface of the brain and coverglass. Blood vessel landmarks were used to find the same fields of view across imaging sessions, and then fine scale adjustments were made to maximize day-to-day overlap by taking short imaging movies (10 seconds) and aligning them to previous days’ recordings.

### Data analysis of imaging and behavioral experiments

#### Extraction of Purkinje cell dendritic ROIs and identification of putative complex spikes

ROIs corresponding to single Purkinje cells were extracted using a combination of Suite2p software in Python (https://github.com/MouseLand/suite2p) (80) for initial source extraction and custom-written software to merge over-segmented dendrites. Procedures were similar to those described previously (22). In general, we found that the updated versions of Suite2p reduced the need for merging of over-segmented dendritic segments.

An event detection algorithm, MLspike (81), was used to identify fast dendritic calcium transients, faithful indicators of complex spiking activity in Purkinje cells (37, 38, 48, 82–84), in each dendritic ROI. Event detection procedures were identical to those described previously, resulting in similar event rates (1.4 ± 0.3 Hz, mean ± s.d.) to our own previous study (22) as well as reported rates of complex spiking during behavior (61). Event amplitudes were converted into rates by multiplying by the imaging frequency (30 Hz), creating a complex spike firing rate weighted by event amplitude.

#### Synchronization of behavior and recordings

All behavioral parameters – trial onset and offset triggers, wheel translation, reward deliveries, tone cues, VR frame update times, 2-photon imaging frame times, and video frame times – were acquired simultaneously and digitized at 5 kHz using a National Instruments card/board (NI USB-6212) and saved using PackIO software (85).

Recorded dendritic fluorescence traces and extracted events (complex spikes) were aligned to different behavioral events of interest at the first frame whose acquisition began after each event and averaged across occurrences of each of these behavioral events. Wheel movement initiation was defined as the first time on each motor trial that wheel velocity exceeded 1 cm/s. Binary licking traces, whose value was one when the mouse’s tongue contacted the lickport and was zero otherwise, were averaged in their raw format in all plots and quantifications.

#### Alignment of PC dendritic segments across recording sessions

Across-day registration of PC dendritic segments was done using a modified version of the Suite2p utility ‘registers2p’ (https://github.com/cortex-lab/Suite2P/tree/master/registers2p). This allows the user to select shared points between 2 reference images (the mean images on any 2 recording days), creating a transformation function that maps one set of image coordinates onto the other image, and warping the ROI coordinates based on this transformation function. Two important modifications were made to the original registers2p code. Firstly, the loading function was amended to take in the Suite2p-python output (after merging over-segmented dendritic segments). Secondly, a manual validation step for each pair of matched ROIs was introduced. Thus, all matched dendritic segments in all paired sessions were manually inspected. For each field of view, the ‘Expert’ session was always used as a reference image, so that ‘Naïve’, ‘Learning’, and ‘Adaptation’ session fields of view were always registered onto the same single reference session. Matched ROIs between non-reference sessions were determined based on transitivity using ‘Expert’ reference sessions.

#### Generalized linear model analysis

To identify task parameters that contributed significantly to the encoding properties of climbing fiber inputs, we fit generalized linear models (GLMs) to the event rates of each PC dendrite based on measured behavioural and task variables using the Matlab version of glmnet (44). A full description of the GLM fitting procedure is provided in **Extended Data Figure 3**.

Behavioural features were downsampled to the acquisition rate of the imaging data (30 Hz), and imaging and behavioural data were trimmed to cover periods from 0.2 s before each trial onset to 1 s after each trial offset. Data from all trials in a session were concatenated to make an initial predictor matrix consisting of time-shifted boxcar filters for discrete events (trial onsets, wheel movement onset, wheel movement offset, reward times, and lick times) and normalized time-shifted versions of continuous variables (object position and wheel velocity). GLMs used to identify significant task encoding (**Figure 3**) were fit on all trials (Undershoot, Correct, and Overshoot) in ‘Expert’ sessions. For each session, we used 2/3 of each trial outcome type as *testing and training trials* and 1/3 of trials in each outcome type (i.e. the second of every set of 3 undershoot, correct, and overshoot trial) as *holdout trials* to test model performance. Thus, testing, training, and holdout trials were taken at approximately equal intervals throughout sessions.

For each PC dendrite, we trained a ‘Gaussian’ GLM using cvglmnet (44) with hyperparameters alpha = 0.5 (elastic-net regularization), nLambda = 100 (to initialize a Lambda sequence that can be optimized), and 10-fold cross-validation. This model was then tested on the *holdout trials* that had not been used for the initial fitting procedure. In order to mitigate the risk of overfitting our training data, we employed an approach based on reduced-rank regression (86) similar to the one used by Steinmetz et al (43). We performed principal component analysis (PCA) on our full predictor matrix and did a parameter sweep across the number of components (ranked by variance explained) to identify the PCA subspace that led to the best model performance on holdout data independently for each Purkinje cell dendrite, creating an optimal predictor matrix (size N components x T timepoints) for each dendrite. The optimal number of components was chosen such that they maximized the deviance explained by the model (calculated as 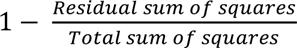, equivalent to R^2^ for Gaussian GLMs). To identify PC dendrites with significant task modulation, we compared the deviance explained by this model to a distribution of null models, which were generated using the same procedure but with the design matrix permuted in time, breaking the relationship between activity and behaviour but maintaining the local structure of each in time (87). Individual PC dendrites were deemed task-modulated if the real deviance explained was (1) above zero, and (2) greater than the mean + 2 standard deviations of the distribution of null models.

To assess if task-modulated PC dendrites were modulated by specific task parameters (e.g. object appearance, movement onset, reward, and licking), we performed the same shuffling procedure described above, but only for the sets of predictors associated with that task parameter. If the full model deviance explained was greater than mean + 2 standard deviations of the distribution of predictor-shuffle models, a given PC dendrite was determined to have significant coding for that task parameter.

For analysis in **Figures 4** and **5**, which required comparable trials across ‘Naïve’, ‘Learning’, ‘Expert’, and ‘Adaptation’ sessions, we re-fit GLMs on recordings from ‘Expert’ sessions as described but using only Correct trials (which were shared across all session types). We could then use recordings from ‘Naïve’, ‘Learning’, and ‘Adaptation’ sessions as the holdout data to test the consistency of task encoding across learning. To determine the mapping between neural activity and either wheel movements or rewards at different task stages, we computed the task feature-specific activity kernel for these task parameters by (1) fitting GLMs with the timing of a given set of predictors shuffled (as we had done previously to determine significant task encoding), (2) subtracting this shuffled prediction from the real activity, leaving us with an activity residual, and (3) refitting a model only using the predictors of interest on the residual activity.

### Statistical analysis

No statistical methods were used to pre-determine sample sizes, but our sample sizes are similar to those reported in previous publications (15, 22). No randomization of experimental subjects was necessary as all mice were trained and recorded under the same conditions. Behavioral events within each training session were randomized on a trial-by-trial basis within the temporal ranges and incidence rates described in the text. Data collection and analysis were not performed blind to the conditions of the experiment, but analysis relied on code that was standardized for all experimental conditions.

Data distributions were not assumed to be normally distributed and all statistical comparisons between groups of continuous variables were performed using non-parametric tests – the Wilcoxon rank-sum test and signed-rank test were used to study differences between two groups of unpaired and paired data, respectively, and the Kruskal-Wallis test and Friedman test were used when comparing more than 2 groups of unpaired and paired data, respectively. Bonferroni correction was applied for multiple comparisons. In general, 95% confidence intervals (p < 0.05) were used to define statistical significance.

## REFERENCES

1. Meister M. Learning, fast and slow. Curr Opin Neurobiol. 2022;75:102555.

2. Liao Z, Losonczy A. Learning, Fast and Slow: Single- and Many-Shot Learning in the Hippocampus. Annu Rev Neurosci. 2024;47(1):187–209.

3. Bourtchuladze R, Frenguelli B, Blendy J, Cioffi D, Schutz G, Silva AJ. Deficient long-term memory in mice with a targeted mutation of the cAMP-responsive element-binding protein. Cell. 1994;79(1):59–68.

4. Haesen K, Beckers T, Baeyens F, Vervliet B. One-trial overshadowing: Evidence for fast specific fear learning in humans. Behav Res Ther. 2017;90:16–24.

5. Krakauer JW, Shadmehr R. Consolidation of motor memory. Trends Neurosci. 2006;29(1):58–64.

6. Shadmehr R, Smith MA, Krakauer JW. Error correction, sensory prediction, and adaptation in motor control. Annu Rev Neurosci. 2010;33:89–108.

7. Peters AJ, Chen SX, Komiyama T. Emergence of reproducible spatiotemporal activity during motor learning. Nature. 2014;510(7504):263–7.

8. Peters AJ, Liu H, Komiyama T. Learning in the Rodent Motor Cortex. Annu Rev Neurosci. 2017;40:77–97.

9. Taylor JA, Ivry RB. The role of strategies in motor learning. Ann N Y Acad Sci. 2012;1251:1–12.

10. Kawai R, Markman T, Poddar R, Ko R, Fantana AL, Dhawale AK, et al. Motor cortex is required for learning but not for executing a motor skill. Neuron. 2015;86(3):800–12.

11. Smith MA, Ghazizadeh A, Shadmehr R. Interacting adaptive processes with different timescales underlie short-term motor learning. PLoS Biol. 2006;4(6):e179.

12. Huberdeau DM, Krakauer JW, Haith AM. Dual-process decomposition in human sensorimotor adaptation. Curr Opin Neurobiol. 2015;33:71–7.

13. Krakauer JW, Hadjiosif AM, Xu J, Wong AL, Haith AM. Motor Learning. Compr Physiol. 2019;9(2):613–63.

14. Kostadinov D, Hausser M. Reward signals in the cerebellum: origins, targets, and functional implications. Neuron. 2022;110:1–14.

15. Wagner MJ, Kim TH, Kadmon J, Nguyen ND, Ganguli S, Schnitzer MJ, et al. Shared Cortex-Cerebellum Dynamics in the Execution and Learning of a Motor Task. Cell. 2019;177(3):669–82 e24.

16. Costa RM, Cohen D, Nicolelis MA. Differential corticostriatal plasticity during fast and slow motor skill learning in mice. Curr Biol. 2004;14(13):1124–34.

17. Gilbert P, Thach W. Purkinje cell activity during motor learning. Brain Research. 1977;128:309–28.

18. Herzfeld DJ, Kojima Y, Soetedjo R, Shadmehr R. Encoding of error and learning to correct that error by the Purkinje cells of the cerebellum. Nat Neurosci. 2018;21(5):736–43.

19. Nguyen V, Gros C, Stell BM. Rapid motor skill adjustment is associated with population-level modulation of cerebellar error signals. Nat Neurosci. 2026;29(1):136–46.

20. Silva NT, Ramirez-Buritica J, Pritchett DL, Carey MR. Climbing fibers provide essential instructive signals for associative learning. Nat Neurosci. 2024;27(5):940–51.

21. De Zeeuw CI. Bidirectional learning in upbound and downbound microzones of the cerebellum. Nat Rev Neurosci. 2021;22(2):92–110.

22. Kostadinov D, Beau M, Blanco-Pozo M, Hausser M. Predictive and reactive reward signals conveyed by climbing fiber inputs to cerebellar Purkinje cells. Nat Neurosci. 2019;22(6):950–62.

23. Ohmae S, Medina JF. Climbing fibers encode a temporal-difference prediction error during cerebellar learning in mice. Nat Neurosci. 2015;18(12):1798–803.

24. Raymond JL, Medina JF. Computational principles of supervised learning in the cerebellum. Annual Review of Neuroscience. 2018;41:233–53.

25. Ikezoe K, Hidaka N, Manita S, Murakami M, Tsutsumi S, Isomura Y, et al. Cerebellar climbing fibers multiplex movement and reward signals during a voluntary movement task in mice. Commun Biol. 2023;6(1):924.

26. Kitazawa S, Kimura T, Yin P-B. Cerebellar complex spikes encode both destinations and errors in arm movements. Nature. 1998;392:494–7.

27. Larry N, Yarkoni M, Lixenberg A, Joshua M. Cerebellar climbing fibers encode expected reward size. Elife. 2019;8.

28. Heffley W, Hull C. Classical conditioning drives learned reward prediction signals in climbing fibers across the lateral cerebellum. Elife. 2019;8.

29. Hull C. Prediction signals in the cerebellum: beyond supervised motor learning. Elife. 2020;9.

30. Welsh JP, Lang EJ, Sugihara I, Llinas R. Dynamic organization of motor control within the olivocerebellar system. Nature. 1995;374:453–7.

31. Simpson JI, Wylie DR, De Zeeuw CI. On climbing fiber signals and their consequence(s). Behavioral and Brain Sciences. 1996;19(03):384–98.

32. Medina JF, Lisberger SG. Links from complex spikes to local plasticity and motor learning in the cerebellum of awake-behaving monkeys. Nat Neurosci. 2008;11(10):1185–92.

33. Medina JF. The multiple roles of Purkinje cells in sensori-motor calibration: to predict, teach and command. Curr Opin Neurobiol. 2011;21(4):616–22.

34. Dhawale AK, Smith MA, Olveczky BP. The Role of Variability in Motor Learning. Annu Rev Neurosci. 2017;40:479–98.

35. Heffley W, Song EY, Xu Z, Taylor BN, Hughes MA, McKinney A, et al. Coordinated cerebellar climbing fiber activity signals learned sensorimotor predictions. Nat Neurosci. 2018;21(10):1431–41.

36. Lee KH, Mathews PJ, Reeves AM, Choe KY, Jami SA, Serrano RE, et al. Circuit mechanisms underlying motor memory formation in the cerebellum. Neuron. 2015;86(2):529–40.

37. Tsutsumi S, Yamazaki M, Miyazaki T, Watanabe M, Sakimura K, Kano M, et al. Structure-function relationships between aldolase C/zebrin II expression and complex spike synchrony in the cerebellum. J Neurosci. 2015;35(2):843–52.

38. Kitamura K, Hausser M. Dendritic calcium signaling triggered by spontaneous and sensory-evoked climbing fiber input to cerebellar Purkinje cells in vivo. J Neurosci. 2011;31(30):10847–58.

39. Gaffield MA, Bonnan A, Christie JM. Conversion of Graded Presynaptic Climbing Fiber Activity into Graded Postsynaptic Ca(2+) Signals by Purkinje Cell Dendrites. Neuron. 2019;102(4):762–9 e4.

40. Pillow JW, Shlens J, Paninski L, Sher A, Litke AM, Chichilnisky EJ, et al. Spatio-temporal correlations and visual signalling in a complete neuronal population. Nature. 2008;454(7207):995–9.

41. Park IM, Meister ML, Huk AC, Pillow JW. Encoding and decoding in parietal cortex during sensorimotor decision-making. Nat Neurosci. 2014;17(10):1395–403.

42. Driscoll LN, Pettit NL, Minderer M, Chettih SN, Harvey CD. Dynamic Reorganization of Neuronal Activity Patterns in Parietal Cortex. Cell. 2017;170(5):986–99 e16.

43. Steinmetz NA, Zatka-Haas P, Carandini M, Harris KD. Distributed coding of choice, action and engagement across the mouse brain. Nature. 2019;576(7786):266–73.

44. Friedman J, Hastie T, Tibshirani R. Regularization Paths for Generalized Linear Models via Coordinate Descent. J Stat Softw. 2010;33(1):1–22.

45. Bayer HM, Glimcher PW. Midbrain dopamine neurons encode a quantitative reward prediction error signal. Neuron. 2005;47(1):129–41.

46. Oscarsson O. Functional units of the cerebellum - sagittal zones and microzones. Trends in Neurosciences. 1979;2:143–5.

47. Lang EJ, Apps R, Bengtsson F, Cerminara NL, De Zeeuw CI, Ebner TJ, et al. The Roles of the Olivocerebellar Pathway in Motor Learning and Motor Control. A Consensus Paper. Cerebellum. 2017;16(1):230–52.

48. Schultz SR, Kitamura K, Post-Uiterweer A, Krupic J, Hausser M. Spatial pattern coding of sensory information by climbing fiber-evoked calcium signals in networks of neighboring cerebellar Purkinje cells. J Neurosci. 2009;29(25):8005–15.

49. Najafi F, Giovannucci A, Wang Samuel SH, Medina Javier F. Sensory-Driven Enhancement of Calcium Signals in Individual Purkinje Cell Dendrites of Awake Mice. Cell Reports. 2014;6(5):792–8.

50. Michikawa T, Yoshida T, Kuroki S, Ishikawa T, Kakei S, Kimizuka R, et al. Distributed sensory coding by cerebellar complex spikes in units of cortical segments. Cell Rep. 2021;37(6):109966.

51. Thach WT. Discharge of cerebellar neurons related to two maintained postures and two prompt movements. II. Purkinje cell output and input. J Neurophysiol. 1970;33(4):537–47.

52. Clothier S, Kostadinov D, Clark BA, Häusser M, Rossi LF. Logic of brain-wide connections with the cerebellar cortex. bioRxiv. 2026.

53. Apps R, Hawkes R. Cerebellar cortical organization: a one-map hypothesis. Nat Rev Neurosci. 2009;10(9):670–81.

54. Bosman LW, Koekkoek SK, Shapiro J, Rijken BF, Zandstra F, van der Ende B, et al. Encoding of whisker input by cerebellar Purkinje cells. J Physiol. 2010;588(Pt 19):3757–83.

55. Lisberger SG. Internal models of eye movement in the floccular complex of the monkey cerebellum. Neuroscience. 2009;162(3):763–76.

56. Soetedjo R, Kojima Y, Fuchs AF. How cerebellar motor learning keeps saccades accurate. J Neurophysiol. 2019;121(6):2153–62.

57. Heiney SA, Wohl MP, Chettih SN, Ruffolo LI, Medina JF. Cerebellar-dependent expression of motor learning during eyeblink conditioning in head-fixed mice. J Neurosci. 2014;34(45):14845–53.

58. Apps R, Hawkes R, Aoki S, Bengtsson F, Brown AM, Chen G, et al. Cerebellar Modules and Their Role as Operational Cerebellar Processing Units: A Consensus paper [corrected]. Cerebellum. 2018;17(5):654–82.

59. Calame DJ, Becker MI, Person AL. Cerebellar associative learning underlies skilled reach adaptation. Nat Neurosci. 2023;26(6):1068–79.

60. Wagner MJ, Savall J, Hernandez O, Mel G, Inan H, Rumyantsev O, et al. A neural circuit state change underlying skilled movements. Cell. 2021;184(14):3731–47 e21.

61. Streng ML, Popa LS, Ebner TJ. Climbing Fibers Control Purkinje Cell Representations of Behavior. J Neurosci. 2017;37(8):1997–2009.

62. Dacre J, Colligan M, Clarke T, Ammer JJ, Schiemann J, Chamosa-Pino V, et al. A cerebellar-thalamocortical pathway drives behavioral context-dependent movement initiation. Neuron. 2021;109(14):2326–38 e8.

63. Sauerbrei BA, Lubenov EV, Siapas AG. Structured Variability in Purkinje Cell Activity during Locomotion. Neuron. 2015;87(4):840–52.

64. Armstrong DM. The supraspinal control of mammalian locomotion. J Physiol. 1988;405:1–37.

65. Hoogland TM, De Gruijl JR, Witter L, Canto CB, De Zeeuw CI. Role of Synchronous Activation of Cerebellar Purkinje Cell Ensembles in Multi-joint Movement Control. Curr Biol. 2015;25(9):1157–65.

66. Beau M, D’Agostino F, Lajko A, Martínez G, Häusser M, Kostadinov D. NeuroPyxels: loading, processing and plotting Neuropixels data in python. Zenodo. 2021.

67. Beau M, Herzfeld DJ, Naveros F, Hemelt ME, D’Agostino F, Oostland M, et al. A deep learning strategy to identify cell types across species from high-density extracellular recordings. Cell. 2025;188(8):2218–34 e22.

68. Steinmetz NA, Aydin C, Lebedeva A, Okun M, Pachitariu M, Bauza M, et al. Neuropixels 2.0: A miniaturized high-density probe for stable, long-term brain recordings. Science. 2021;372(6539).

69. Li N, Mrsic-Flogel TD. Cortico-cerebellar interactions during goal-directed behavior. Curr Opin Neurobiol. 2020;65:27–37.

70. Agueci L, Cayco-Gajic NA. Distributed learning across fast and slow neural systems supports efficient motor adaptation. bioRxiv. 2025.

71. French RM. Catastrophic forgetting in connectionist networks. Trends Cogn Sci. 1999;3(4):128–35.

72. Kirkpatrick J, Pascanu R, Rabinowitz N, Veness J, Desjardins G, Rusu AA, et al. Overcoming catastrophic forgetting in neural networks. Proc Natl Acad Sci U S A. 2017;114(13):3521–6.

73. Dohare S, Hernandez-Garcia JF, Lan Q, Rahman P, Mahmood AR, Sutton RS. Loss of plasticity in deep continual learning. Nature. 2024;632(8026):768–74.

74. Zhang XM, Ng AH, Tanner JA, Wu WT, Copeland NG, Jenkins NA, et al. Highly restricted expression of Cre recombinase in cerebellar Purkinje cells. Genesis. 2004;40(1):45–51.

75. Chen TW, Wardill TJ, Sun Y, Pulver SR, Renninger SL, Baohan A, et al. Ultrasensitive fluorescent proteins for imaging neuronal activity. Nature. 2013;499(7458):295–300.

76. Dana H, Sun Y, Mohar B, Hulse BK, Kerlin AM, Hasseman JP, et al. High-performance calcium sensors for imaging activity in neuronal populations and microcompartments. Nat Methods. 2019;16(7):649–57.

77. Aronov D, Tank DW. Engagement of neural circuits underlying 2D spatial navigation in a rodent virtual reality system. Neuron. 2014;84(2):442–56.

78. Slotnick B. A simple 2-transistor touch or lick detector circuit. J Exp Anal Behav. 2009;91(2):253–5.

79. Pologruto TA, Sabatini BL, Svoboda K. ScanImage: Flexible software for operating laser scanning microscopes. BioMedical Engineering OnLine. 2003;2(13).

80. Pachitariu M, Stringer C, Dipoppa M, Schröder S, Rossi LF, Dalgleish H, et al. Suite2p: beyond 10,000 neurons with standard two-photon microscopy. bioRxiv. 2017.

81. Deneux T, Kaszas A, Szalay G, Katona G, Lakner T, Grinvald A, et al. Accurate spike estimation from noisy calcium signals for ultrafast three-dimensional imaging of large neuronal populations in vivo. Nat Commun. 2016;7:12190.

82. Ozden I, Lee HM, Sullivan MR, Wang SS. Identification and clustering of event patterns from in vivo multiphoton optical recordings of neuronal ensembles. J Neurophysiol. 2008;100(1):495–503.

83. Ozden I, Sullivan MR, Lee HM, Wang SS. Reliable coding emerges from coactivation of climbing fibers in microbands of cerebellar Purkinje neurons. J Neurosci. 2009;29(34):10463–73.

84. Ramirez JE, Stell BM. Calcium Imaging Reveals Coordinated Simple Spike Pauses in Populations of Cerebellar Purkinje Cells. Cell Rep. 2016;17(12):3125–32.

85. Watson BO, Yuste R, Packer AM. PackIO and EphysViewer: software tools for acquisition and analysis of neuroscience data. bioRxiv. 2016.

86. Keeley SL, Zoltowski DM, Aoi MC, Pillow JW. Modeling statistical dependencies in multi-region spike train data. Curr Opin Neurobiol. 2020;65:194–202.

87. Harris KD. Nonsense correlations in neuroscience. bioRxiv. 2021.

