## Extended Data for "Fast and slow learning mediated by distinct climbing fiber signals"

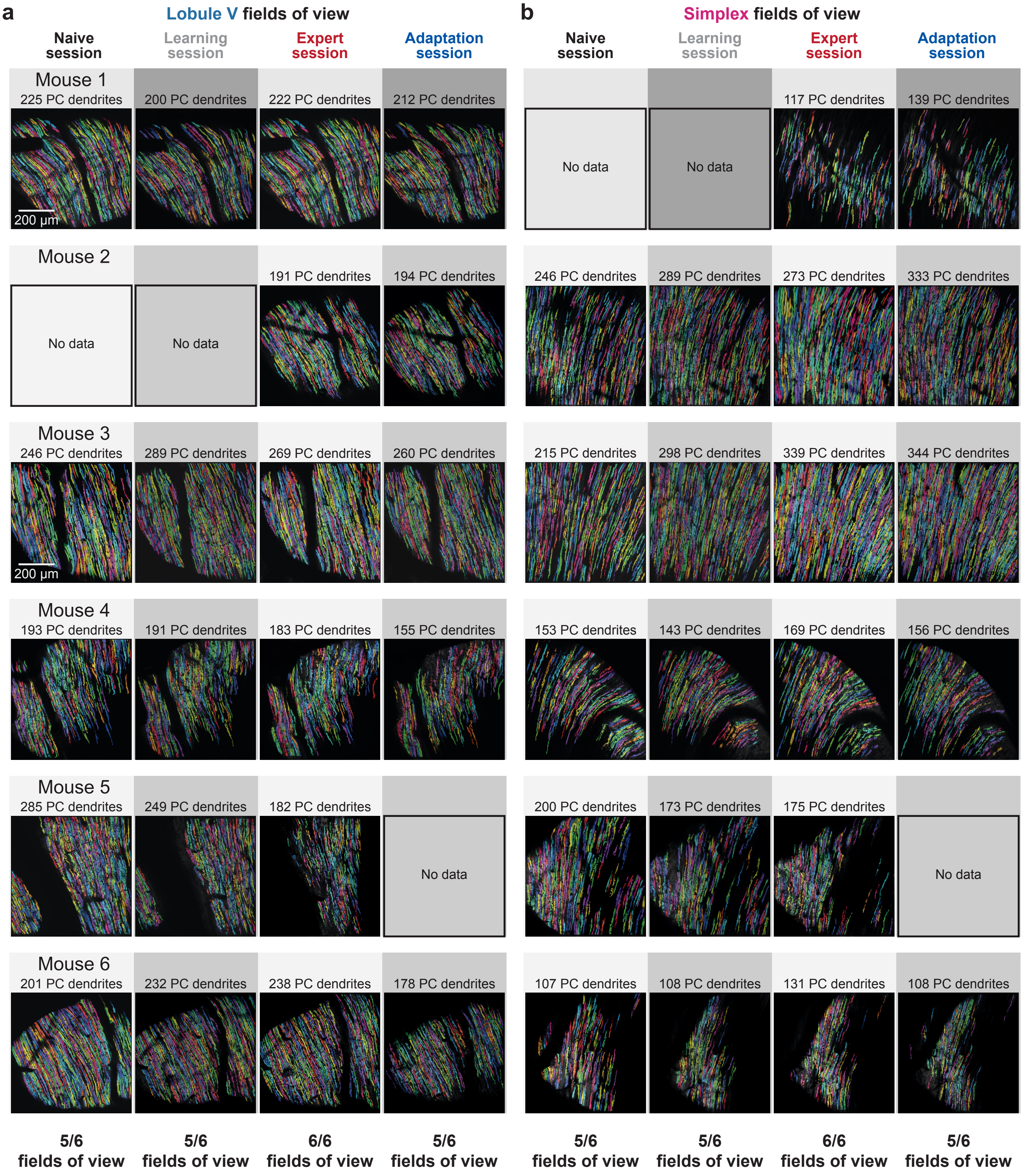


**Extended Data Figure 1**: All chronically recorded fields of view

**EXTENDED DATA FIGURE 1: All chronically recorded fields of view**

Chronically recorded fields of view in both Lobule V (panel **a**) and Lobule simplex (panel **b**) are shown for all mice recorded in this study. Not all mice have data in all fields of view across all session types, but both Lobule V and simplex were recorded in at least 5 of 6 mice at each task stage.

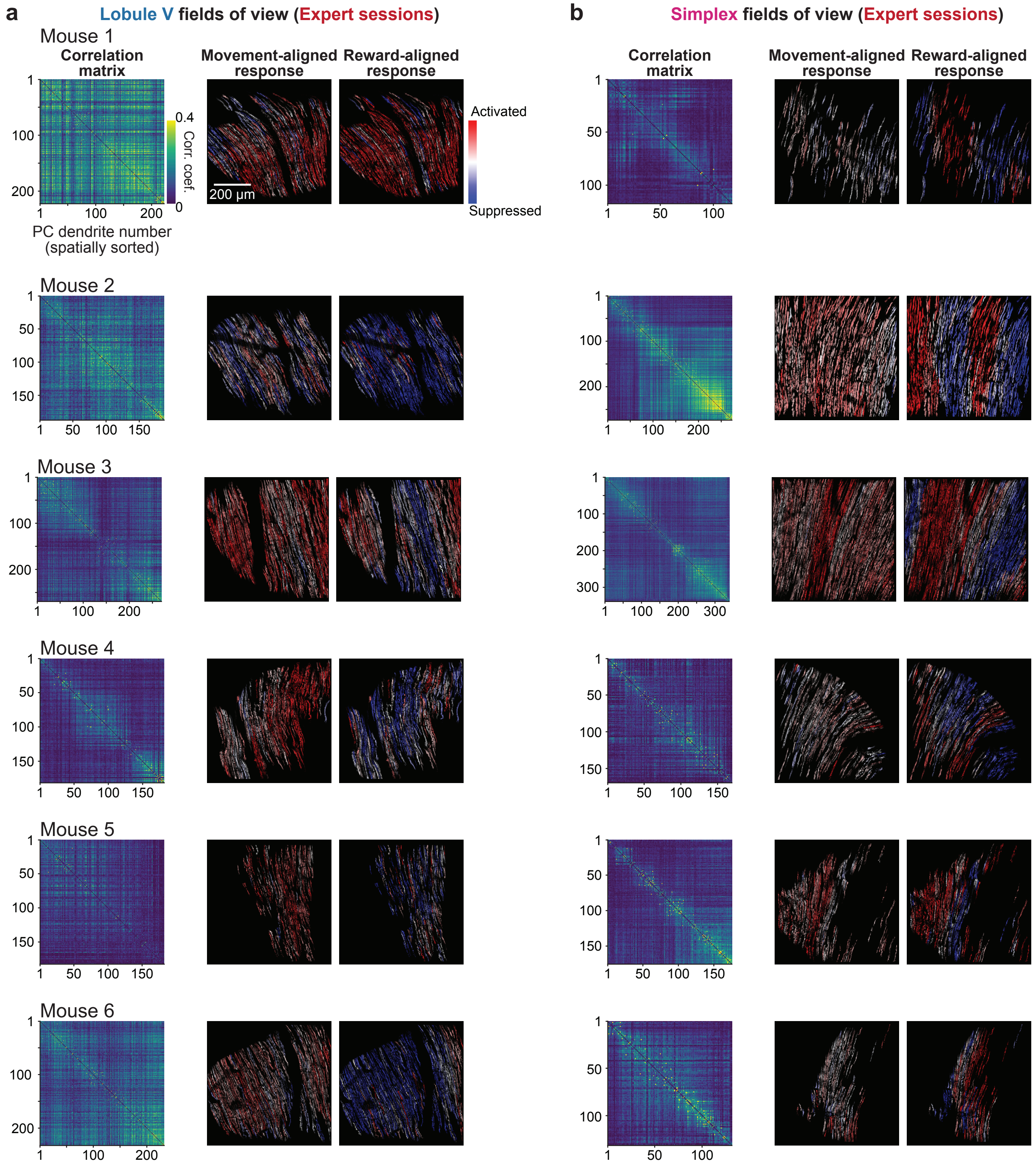


**Extended Data Figure 2**: Spatial organization of movement and reward-aligned responses

**EXTENDED DATA FIGURE 2:**

**Spatial organization of movement and reward-aligned responses**

**a.** Left panels depict the correlation matrices of fluorescence activity recorded in spatially sorted (medial to lateral) PC dendrites in Lobule V for each field of view. Middle and right panels show the movement-aligned response selectivity (computed as mean activity over -300 to 0 ms from movement onset) and reward-aligned response selectivity (computed as mean activity over 0 to +200 ms from reward delivery) for each field of view. Selectivity of each PC dendrite was calculated as the log of baseline-normalized activity for each task event.

**b.** Same as panel **a** but for Lobule simplex fields of view.

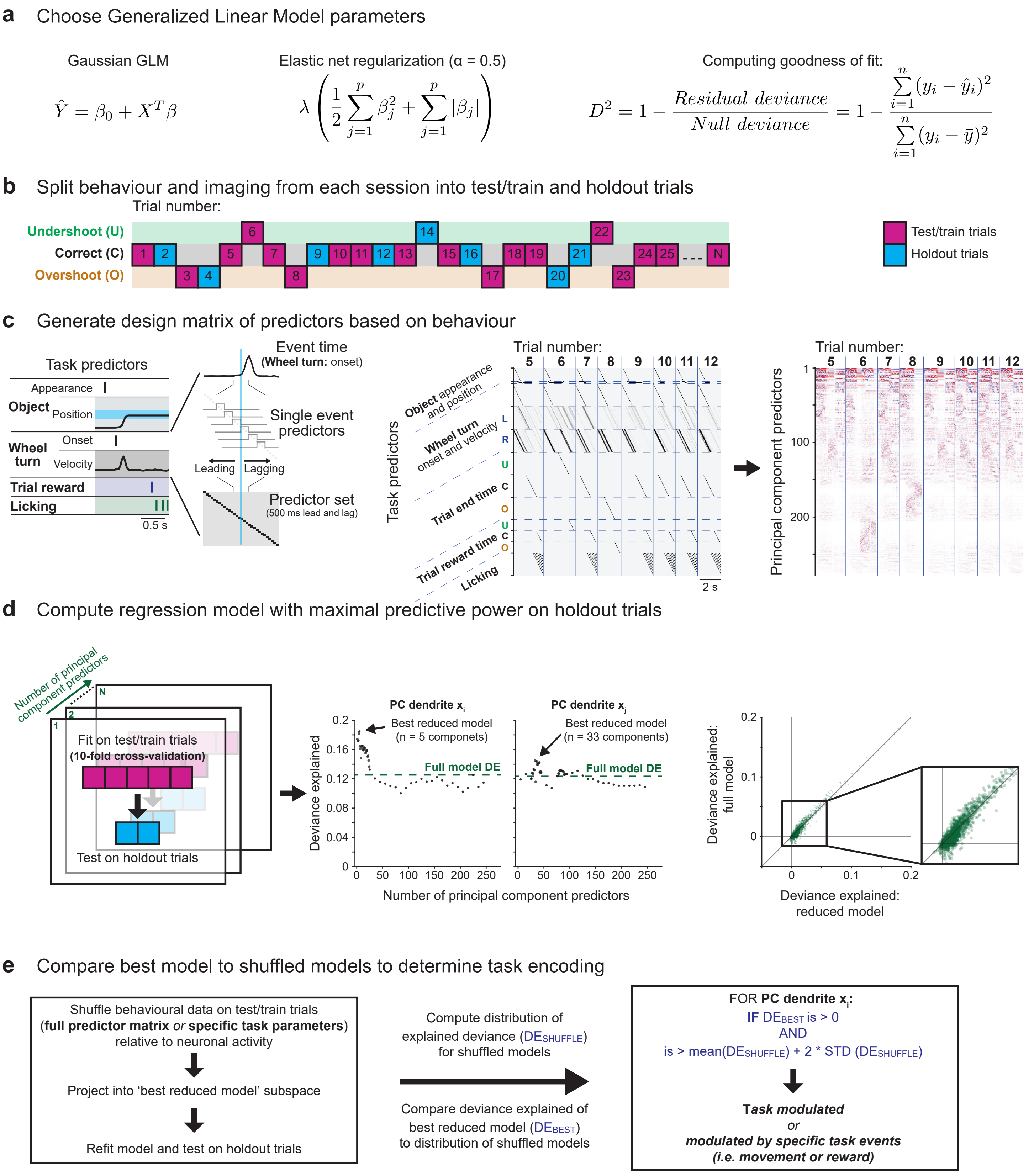


**Extended Data Figure 3**: Generalized linear model fitting procedures

**EXTENDED DATA FIGURE 3: Generalized linear model fitting procedures**

**a.** Choice of Gaussian GLM, Elastic net regularization with equal weighting of lasso and ridge regularization, and Deviance explained formation is depicted.

**b.** Visualization of data partitioning into test/train and holdout trials for each session: For each trial outcome type (undershoot, correct, and overshoot), we split the trials into consecutive groups of 3. We used the first and third trial of each set to test and train our models and the second trial as holdout data to assess model performance.

**c.** Generation of design matrix: After downsampling our behaviour data to our imaging rate (30 Hz), behavioural data from all trials in a session were concatenated to make an initial predictor matrix consisting of time-shifted boxcar filters for discrete events (trial onsets, wheel movement onset, wheel movement offset, reward times, and lick times) and normalized time-shifted versions of continuous variables (object position and wheel velocity). We then performed principal component analysis (PCA) on our full predictor matrix, which we could use to mitigate overfitting of our training data.

**d.** Depiction of approach to identify the model with maximal predictive power: We performed a parameter sweep of our PCA predictor matrix to identify the number of components (rows of the PCA matrix sorted by variance explained) that yielded the best deviance explained ($DE_{BEST}$ model) on our heldout trials (left and middle). This approach yielded an improvement of deviance explained across all recorded neurons when compared to a model using our initial, full predictor matrix (right).

**e.** Comparison to null models: To identify which neurons were significantly task-modulated or modulated by specific task events, we employed a shuffling procedure in which we circularly permuted our behaviour data in time (all predictors together for general task modulation or specific subsets of predictors for task event modulation). We then projected this matrix into the PCA subspace used for the $DE_{BEST}$ model, and refit on the holdout trials. Repeating this procedure yielded a distribution of shuffle models ($DE_{SHUFFLE}$), which we could compare against $DE_{BEST}$ to assess statistically significant task modulation.

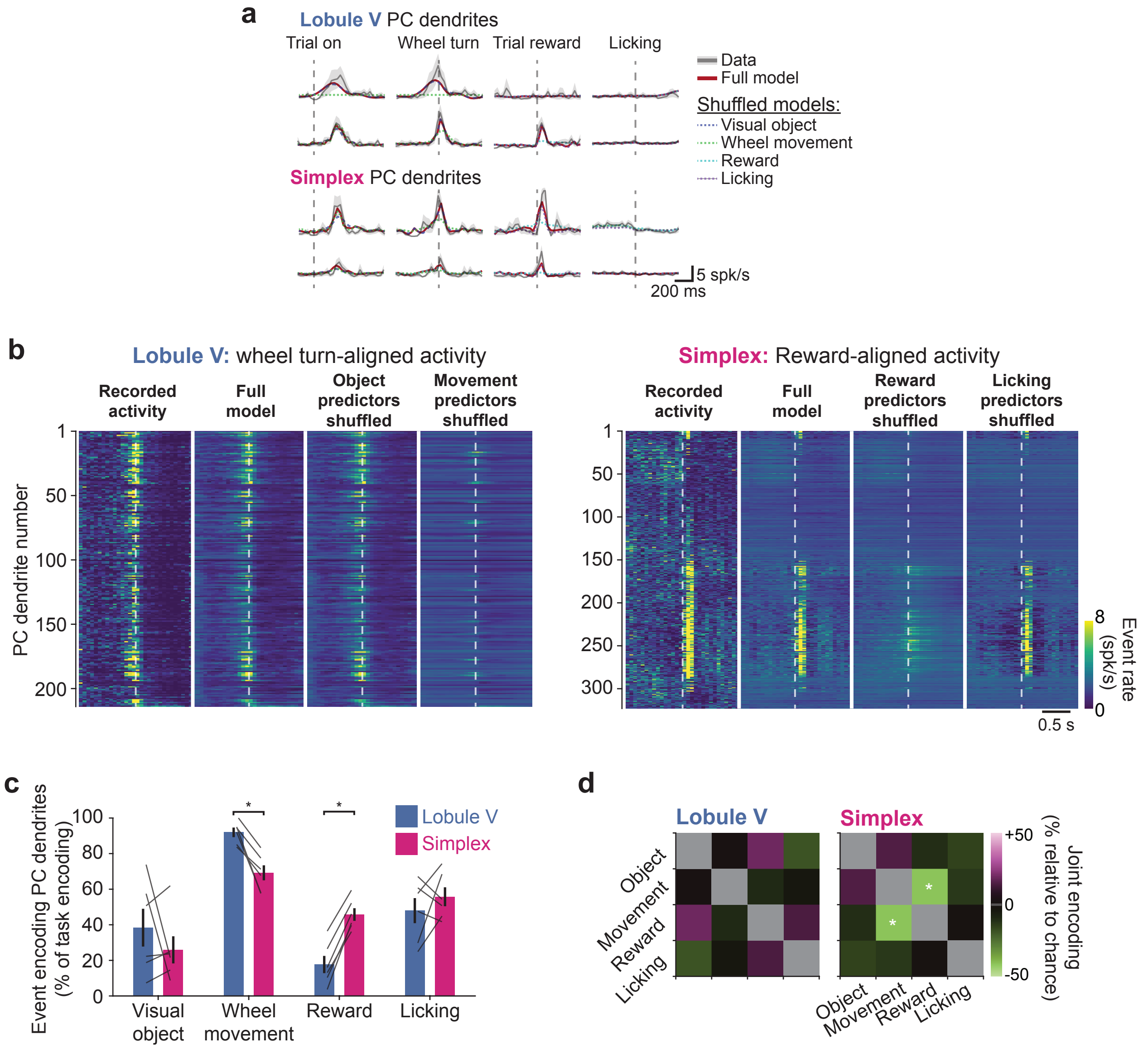


**Extended Data Figure 4**: Performance of generalized linear model across mice and regions

**EXTENDED DATA FIGURE 4:**
**Performance of generalized linear models across mice and regions**

**a.** Example PC dendritic activity aligned to trial onset, wheel turns, rewards, and licking. Individual traces show real data, full best model fit, and model fits with specific task event parameters shuffled.

**b.** Task event-aligned activity for full fields of view in Lobule V and simplex, shown as heatmaps, allowing for visualization of differences in task encoding when specific task event parameters are shuffled.

**c.** Percentage of task-encoding PC dendrites that encode specific task events in expert mice (n = 6 sessions in 6 mice, 1 session per field of view per mouse). Statistics: Lobule V vs simplex – P = 1 [Visual object]; P = 0.031 [Wheel movement]; P = 0.031 [Reward]; P = 0.44 [Lick]; Wilcoxon signed-rank test.

**d.** Likelihood of joint encoding of multiple task events, shown as a comparison to expected joint encoding by chance i.e. the product of independent probabilities (n = 6 sessions each for Lobule V and simplex from 6 mice, 1 session per region per mouse). Statistical comparisons – Lobule V: P = 0.977 [Object + Movement], P = 0.416 [Object + Reward]; P = 0.333 [Object + Licking]; P = 0.21 [Movement + Reward]; P = 0.201 [Movement + Licking]; P = 0.911 [Reward + Licking]. Statistical comparisons – Lobule simplex: P = 0.150 [Object + Movement], P = 0.252 [Object + Reward]; P = 0.437 [Object + Licking]; P = 0.035 [Movement + Reward]; P = 0.186 [Movement + Licking]; P = 0.995 [Reward + Licking].

Data are shown as mean ± s.e.m. unless otherwise noted. Statistical tests: $*P < 0.05$.

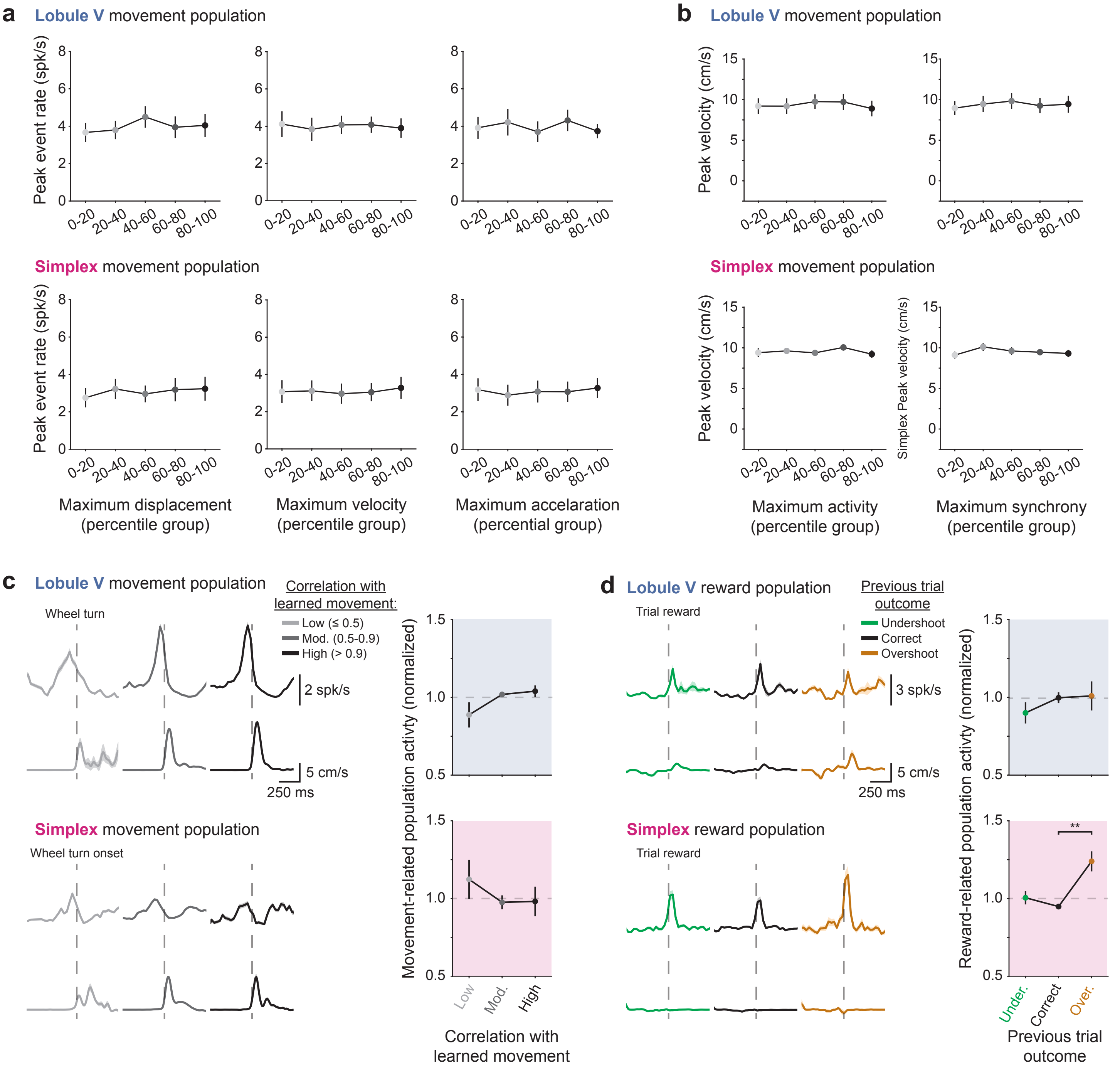


**Extended Data Figure 5**: Kinematic and performance-based encoding by climbing fibers

**EXTENDED DATA FIGURE 5: Kinematic and performance-based encoding by climbing fibers**

**a.** The average peak event rate per dendrite in movement-coding PC dendrites in Lobule V (top) and simplex (bottom) is plotted as a function of wheel displacement, velocity, and acceleration. Data groups on the x-axis are binned into quintiles.

**b.** Peak wheel velocity per trial is plotted as a function of maximum activity and synchrony levels of movement-coding PC dendrites in Lobule V (top) and simplex (bottom). Data groups on the x-axis are binned into quintiles.

**c.** **Left:** Example movement-aligned activity in movement-coding PC dendrite populations in Lobule V (top) and simplex (bottom) in single sessions, split based on the correlation of individual movements to the learned movement. **Right:** Summary of peak movement-related population activity for all recorded mice (n = 6 sessions in 6 mice, 1 session per field of view per mouse). Statistics: Lobule V – Friedman test, Chi-squared(2) = 1.33, P = 0.514 (non-significant); Lobule simplex – Friedman test, Chi-squared(2) = 1.33, P = 0.514 (non-significant).

**d.** **Left:** Example reward-aligned activity in reward-coding PC dendrite populations in Lobule V (top) and simplex (bottom) in single sessions, split based on the outcome of the previous trial. **Right:** Summary of peak reward-related population activity for all recorded mice (n = 6 sessions in 6 mice, 1 session per field of view per mouse). Statistics: Lobule V – Friedman test, Chi-squared(2) = 1.00, P = 0.607 (non-significant); Lobule simplex – Friedman test, Chi-squared(2) = 10.33, P = 0.006, significance values for Bonferroni-corrected individual comparisons: previous trial undershoot vs correct, P = 0.745; previous trial undershoot vs overshoot, P = 0.13; previous trial correct vs overshoot, P = 0.005.

Data are shown as mean ± s.e.m. unless otherwise noted. Statistical tests: ***P < 0.01*.

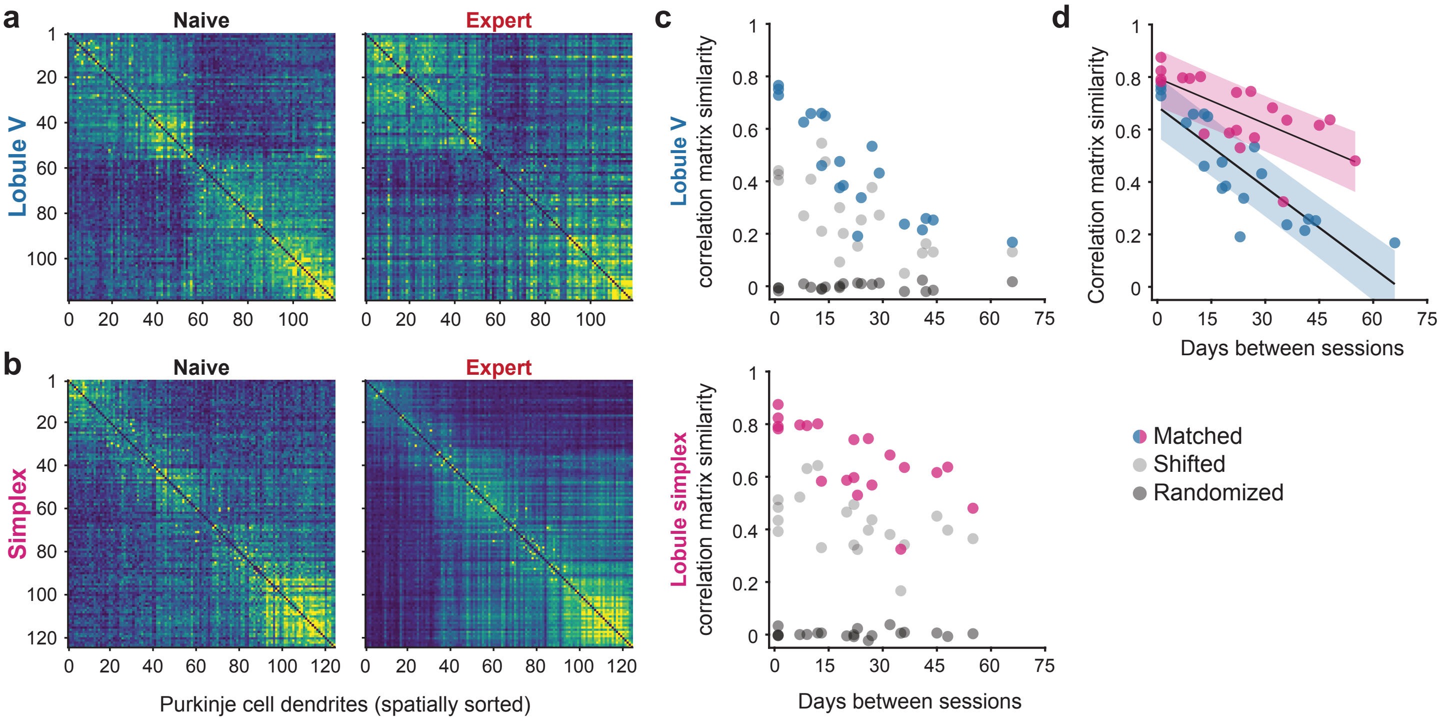


**Extended Data Figure 6**: Correlation structure of climbing fiber inputs across learning

**EXTENDED DATA FIGURE 6: Correlation structure of climbing fiber inputs across learning**

- **a.** Example correlation heatmaps (spatially sorted) of PC dendrites in Lobule V matched across a Naïve and Expert session.
- **b.** Same as panel **a** for a Lobule simplex field of view.
- **c.** Scatterplot of correlation matrix similarity as a function of days between sessions for PC dendrites in Lobule V (blue, top) and simplex (magenta, bottom). Real correlation matrix similarities are shown in color and compared to shifted matrix versions that compare PC dendrites to their nearest neighbours' activity (gray) as well as PC dendrites with locations randomly shuffled (black).
- **d.** Linear fits comparing correlation matrix similarity in Lobules V and simplex as a function of days between recording sessions.

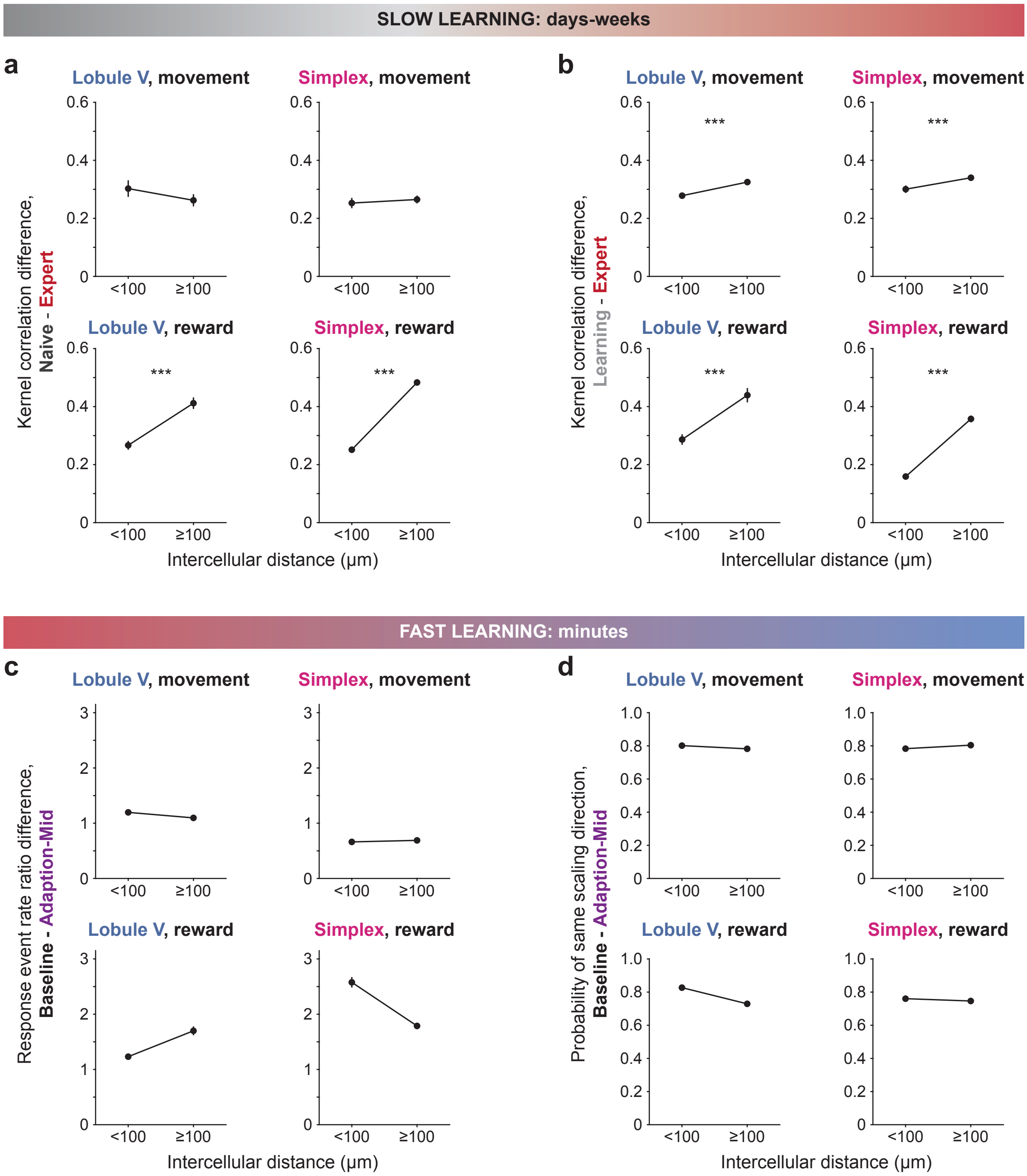


**Extended Data Figure 7**: Distance-dependence of learning-related changes

**EXTENDED DATA FIGURE 7: Distance-dependence of learning-related changes**

**a.** Comparison of the difference between kernel correlations across Naïve and Expert sessions in PC dendrites encoding task events (movement, reward) that are less than or greater than or equal to 100 μm apart. Statistics: Lobule V (Reward), P = 2.6 x $10^{-6}$ (n = 101 comparisons [<100 μm] and n = 193 [≥100 μm]); Lobule simplex (Reward), P = 2.6 x $10^{-48}$ (n = 740 [<100 μm] and n = 1107 [≥100 μm]), Wilcoxon rank-sum test. Data from individual PC dendrites pooled from 5 mice.

**b.** Same as panel **a** for Learning and Expert sessions. Statistics: Lobule V (Movement), P = 2.2 x $10^{-8}$ (n = 1881 comparisons [<100 μm] and n = 3240 [≥100 μm]); Lobule V (Reward), P = 1.4 x $10^{-4}$ (n = 90 [<100 μm] and n = 184 [≥100 μm]); Lobule simplex (Movement), P = 0.0097 (n = 358 [<100 μm] and n = 591 [≥100 μm]); Lobule simplex (Reward), P = 2.4 x $10^{-30}$ (n = 762 [<100 μm] and n = 1211 [≥100 μm]), Wilcoxon rank-sum test. Data from individual PC dendrites pooled from 5 mice.

**c.** Comparison of the difference between adaptation ratios (computed as the responses during Adaptation-mid trials divided by the responses during baseline trials in Adaptation sessions) in movement and reward-encoding PC dendrites.

**d.** Comparison of likelihood of similar Adaptation session scaling direction (both increasing or both decreasing) for movement and reward-encoding PC dendrites.

Data are shown as mean ± s.e.m. unless otherwise noted. Statistical tests: ****P < 0.001*.

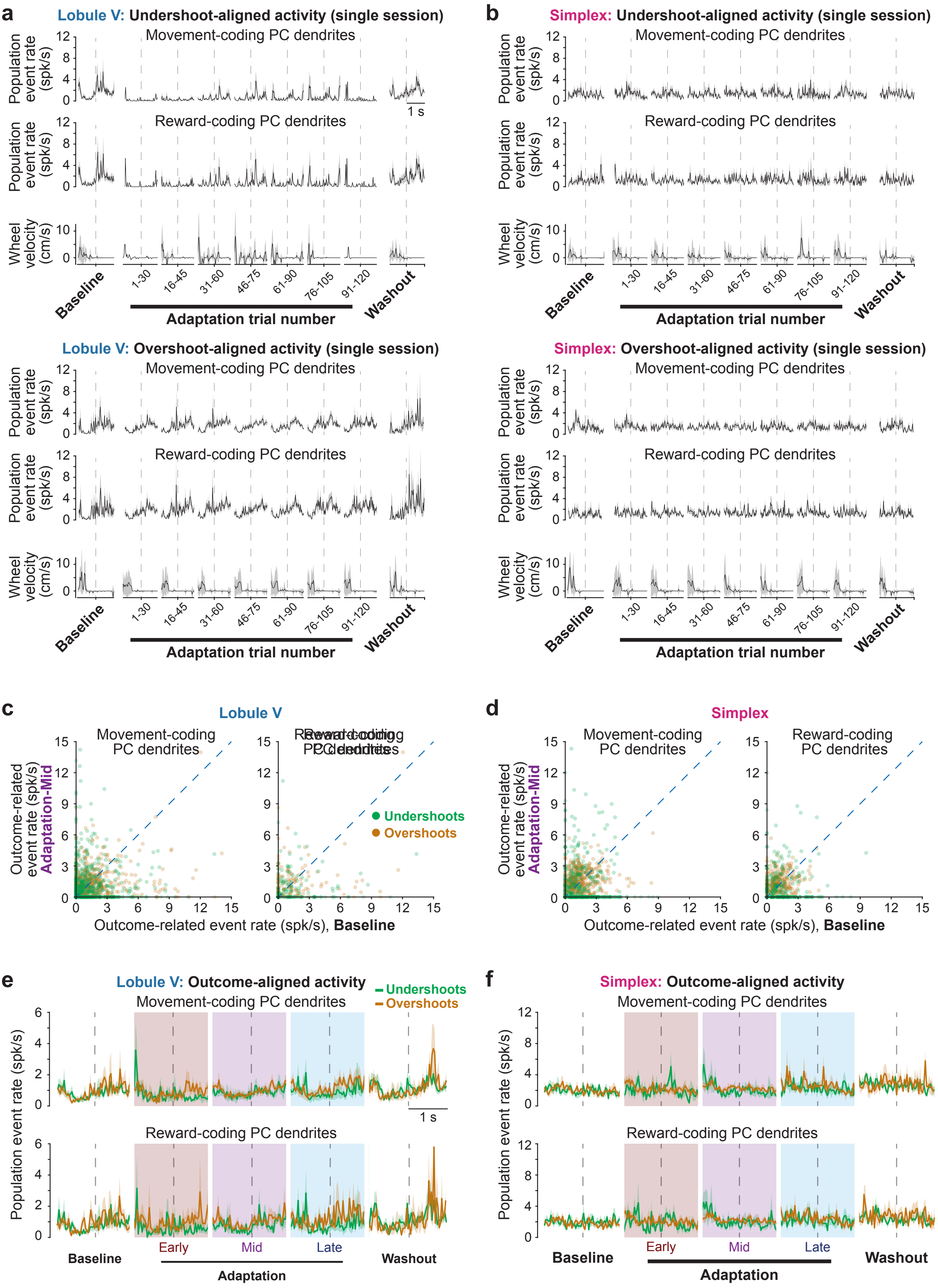


**Extended Data Figure 8**: Climbing fiber dynamics on incorrect trials during adaptation

**EXTENDED DATA FIGURE 8:**

**Climbing fiber dynamics on incorrect trials during adaptation**

**a.** Incorrect outcome-aligned activity of task-encoding PC dendrites in Lobule V of an example mouse in a single Adaptation session. Data are shown in groups of 30 trials that overlap by 15 trials. **Top**: Mean time course of dendritic event rate aligned to expected reward time (dashed line) on undershoot trials in movement-coding (n = 132) and reward-coding (n = 55) PC dendrites. **Bottom**: Mean time course of dendritic event rate aligned to expected reward time (dashed line) on overshoot trials in the same PC dendrites.

**b.** Same as panel **a** but for example Lobule simplex recording session (n = 36 movement-coding and n = 34 reward-coding PC dendrites).

**c.** Scatter plots showing pairwise comparisons of incorrect outcome-related activity during baseline trials and mid-adaptation trials in movement- and reward-encoding PC dendrites in Lobule V (n = 602 [movement] and 128 [reward] dendrites from 5 mice). Undershoot and overshoot responses are shown in green and orange, respectively. Responses relative to task events are computed as in **Figure 5**.

**d.** Same as panel **c** but for Lobule simplex recording sessions (n = 413 [movement] and 274 [reward] dendrites from 5 mice) during baseline trials and mid-adaptation trials (computed as mean activity over 0 to +200 ms from reward delivery).

**e.** Average incorrect outcome-aligned activity in Lobule V movement-coding (top) and reward-coding PC dendrites for baseline, adaptation (early, mid, and late), and washout trials (n = 5 sessions in 5 mice, 1 session per field of view per mouse).

Same as panel **e** but for Lobule simplex (n = 5 sessions in 5 mice, 1 session per field of view per mouse).
